# Fluor NMR study of amino acid derived ligand to study TSPO

**DOI:** 10.1101/2024.10.14.618300

**Authors:** Luminita Duma, Severine Schneider, Agathe Martinez, Cathy Hachet, Frederic Bihel, Jean-Jacques Lacapere

## Abstract

Translocator protein (TSPO, 18 kDa), previously known as peripheral-type benzodiazepine receptor, is an evolutionarily conserved membrane protein involved in various physiological processes and patho-physiological conditions. The endogeneous TSPO ligand is a polypeptide of 9 kDa, but dipeptides with biological activity have been previously synthesized and characterized. Herein, we synthesized a phenyl alanine derived ligand with a ^19^F labelling which opens prospective for ^19^F-MRI and potential ^18^F-PET applications. We characterized the coexistence of two conformers that are not equally sensitive to the media used for membrane protein studies. Interaction studies with the recombinant mouse TSPO (mTSPO) in different membrane-mimicking environments are presented using ^19^F NMR enabling structure/function characterizations. A change in the mTSPO environment from pure detergent to lipid/detergent mixture reveals different exchange rates between bound and free ligand forms. Competition experiments with the high-affinity drug ligand (*R*)-PK 11195 suggests that phenyl alanine derived ligand binds in the same protein cavity.

**Highlights:**

- Fluor labelling of ligands easily reveals the presence of conformers
- Fluorinated phenyl alanine derived ligand interacts with TSPO
- Fluor NMR enables characterization of TSPO ligand interactions
- Fluor NMR facilitates exchange rate studies between free and bound ligand states

**Graphical Abstract:** 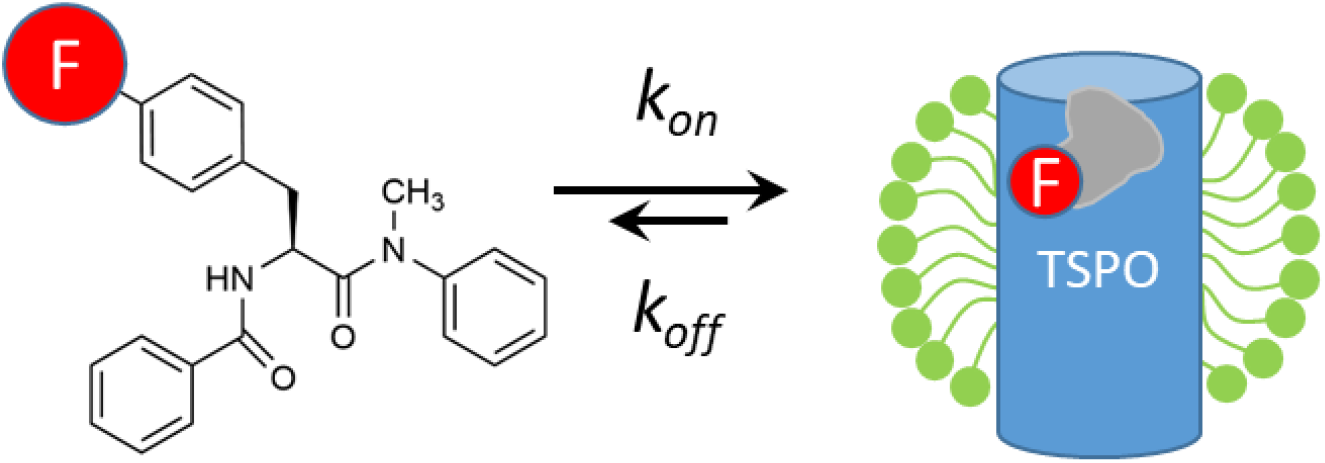

## 1. Introduction

TSPO, previously named PBR for Peripheral-type Benzodiazepine Receptor, was identified in a search for diazepam-binding site [1]. Later on, a polypeptide of 9 kDa was characterized as a diazepam-binding inhibitor, DBI [2] that binds to TSPO with multiple biological actions. More recently, dipeptides active on TSPO but without the side effects of benzodiazepine tranquilizers, were synthesized and showed anxiolytic effects [3–7]. However, no atomic structure of any of these TSPO-peptide complexes is available yet and the mechanism of action remains unknown.

Since TSPO overexpression has been observed in various diseases associated with inflammatory or malignant tumours, TSPO specific ligands have been used as diagnostic and therapeutic agents [8,9]. Different classes of ligands have been developed for Positron Emission Tomography (PET) imaging [10– 13]. Among them, ^11^C and ^18^F derivatives of non-benzodiazepine-type with high affinity for TSPO (PK 11195 and PK 14105, respectively) have been synthesized [14,15] and used in the early development of PET scan imaging. Compared to the first developed ^11^C radiolabelled TSPO ligands used in PET scan, the ^18^F radiolabelled tracers are characterized by a longer lifetime (t_1/2_ = 109.8 min (^18^F) vs 20.3 min (^11^C)), property particularly advantageous for clinical applications. More recently, ^19^F labelled molecules have been successfully used for multimodal ^1^H/^19^F Magnetic Resonance Imaging (MRI) combined with optical fluorescence imaging, various tomography and acoustic imaging techniques. Interestingly, the weak endogenous ^19^F signal in the body favours ^19^F MRI in terms of sensitivity or accuracy [16,17].

At a molecular level, a ^19^F-labelled ligand enables to overcome the overlap between ligand and protein signals that are present in ^1^H Nuclear Magnetic Resonance (NMR) spectra, and prevents the protein isotope labelling. ^19^F ligands are particularly interesting in ^19^F NMR spectroscopy to probe protein-ligand interactions in classical or competition titrations [18] especially for ligands interacting with membrane proteins, which are often purified in detergent and even lipids-containing buffers [19–22].

In the present study, we synthesized a new amino acid derived fluorinated ligand, and characterized by ^19^F NMR spectroscopy its binding to the mouse translocator protein (mTSPO). In this proof-of-concept work on the ligand-membrane protein, with a focus on mTSPO, we explored the effect of the membrane protein environment upon the affinity of the interaction.

## 2. Materials and methods

### 2.1 Synthesis and compounds characterization

**Scheme 1:**
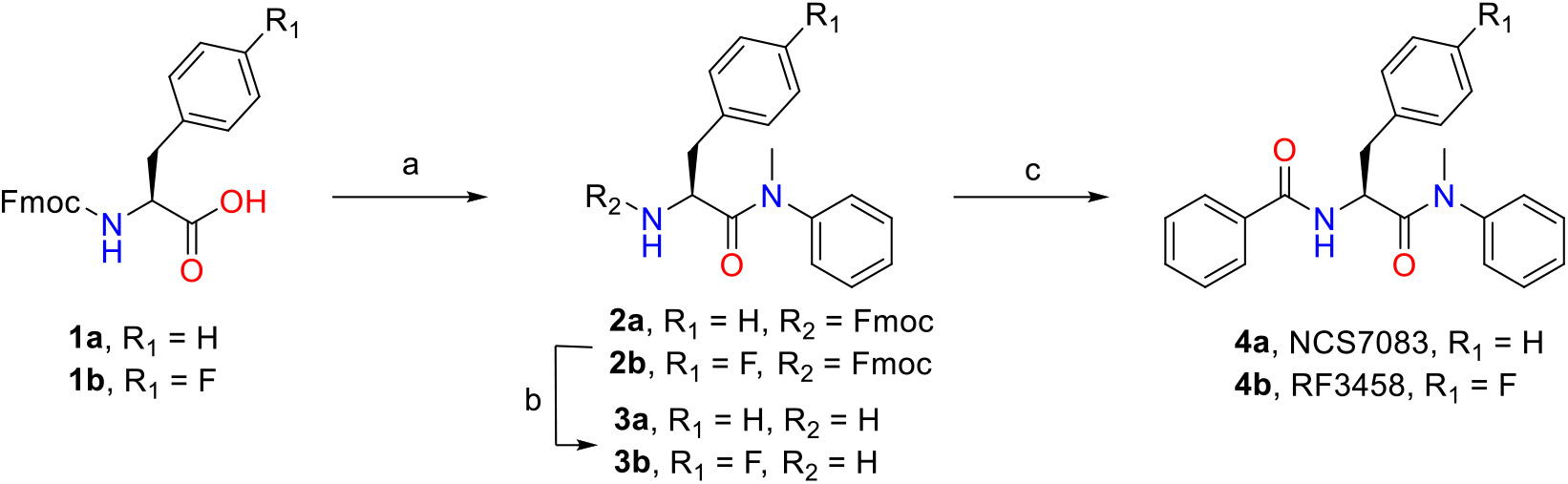
Synthesis of NCS7083 (4a) and RF3458 (4b). Reagents and conditions: a) N-Me-aniline, HATU, DIEA, DCM:DMF, room temperature; b) DBU, DCM; c) BzCl, DIEA, DCM, room temperature.

General procedure for the synthesis of NCS7083 (**4a**) and RF3458 (**4b**): Commercially available Fmoc-L-Phe-OH **1a** (0.37 mmol) or Fmoc-L-Phe(4F)-OH **1b** (0.37 mmol) and HATU (0.39 mmol) were dissolved in a mixture of DCM and DMF (2.5 mL and 0.6 mL, respectively). DIEA (1.1 mmol) was added and the mixture was stirred at room temperature for 10 minutes. N-Methylaniline (0.44 mmol) was then added and the mixture was stirred overnight at room temperature. After evaporation under vacuum, the crude residue was dissolved in EtOAc (10 mL) and successively washed with 1 M aqueous HCl (3 × 10 mL), saturated aqueous NaHCO_3_ (3 × 10 mL), and brine (3 × 10 mL). After evaporation under vacuum, the crude product was purified by flash chromatography on silica gel using a mixture of DCM and EtOAc as the eluent. The purified products **2a** and **2b** were obtained as white powders in 67% and 61% yield, respectively.

Compound **2a** or **2b** (0.23 mmol) was dissolved in DCM (2.5 mL), followed by DBU (0.45 mmol), and the reaction mixture was stirred at room temperature. After reaction completion, the reaction mixture was directly purified on silica gel using DCM:MeOH as the eluent. Compounds **3a** and **3b** (0.2 mmol) was recovered as a yellow oil in 88% and 89% yield, respectively. Compounds **3a** and **3b** were directly used in the next step and dissolved in DCM (2.2 mL), followed by DIEA (0.6 mmol) and benzoyl chloride (0.21 mmol). The reaction mixture was stirred at room temperature until completion. After evaporation under vacuum, the crude residue was dissolved in EtOAc (10 mL) and successively washed with 1 M aqueous HCl (3 × 10 mL), saturated aqueous NaHCO_3_ (3 × 10 mL), and brine (3 × 10 mL). After evaporation under vacuum, the crude product was purified by flash chromatography on silica gel using a mixture of DCM and EtOAc as the eluent. ^1^H and ^13^C NMR chemical shifts of **4a** (NCS7083) are in agreement with the ones reported previously [23]. ^1^H and ^13^C spectra of **4a** (NCS7083) in chloroform and the chromatogram are shown in Figure S1. ^1^H. ^13^C and ^19^F spectra of RF3458 in chloroform are presented in Figure S2. All chemical shifts of **4a** (NCSF7083) and **4b** (RF3458) are summarized in Table S1. The purified products **4a** (NCS7083) and **4b** (RF3458) were obtained as white powders in 93% and 66% yield, respectively. The NMR assignment of the final products is presented in the supplementary material.

### 2.2 Absorbance and solubility screening

Absorbance of fluorinated (RF3458) and non-fluorinated (NCS7083) compounds, respectively, has been recorded on a Carry 60 UV-Vis (Agilent, les Ullis, France). The solubility in ethanol has been obtained by trials and errors and checked by absorbance to be around ∼14 mM. The UV spectrum of RF3458 reveals a small peak at 272 nm and a massif at 226 nm, which permits to calculate the molar extinction coefficient of 170 and 9400 M^-1^cm^-1^, respectively in ethanol (Figure S3A and B). Conversely, the non-fluorinated NCF7083 does not exhibit a specific absorption peak but only shows a massif at 230 nm leading to the determination of a molar extinction coefficient of 25250 M^-1^cm^-1^ in ethanol (data not shown). The solubility in phosphate buffer in the absence and in the presence of detergents has also been measured following the absorbance at 250 and 300 nm and the light scattering induced by the non-soluble compound, respectively (Figure S3C and D). The presence of detergent increases the solubility: therefore, it goes from ∼0.2 mM (without detergent) to ∼0.4 mM (in the presence of 0.1% DPC), respectively. It has to be noted that the solubility increases with the increase of the detergent concentration.

### 2.3 NMR experiments

Experiments have been recorded on Bruker Avance III or Neo spectrometers operating at 600, 500 or 400 MHz for ^1^H, 150, 126 or 101 MHz for ^13^C, and 470 MHz for ^19^F. ^1^H, ^13^C and ^19^F NMR spectra (Figure S2) at room temperature in CDCl_3_ were recorded for the RF3458 (∼10 mg/mL). To see the effect of concentration on ^19^F signals, ^19^F spectra have been recorded at 5, 0.5 and 0.05 mM RF3458 in EtOH, 20% D_2_O (Figure 2A). RF3458 (0.038 µg/mL or 100 µM) has been also characterized in 40 mM phosphate buffer pH 6.5, DPC 0.2%, 10% D_2_O, containing as internal standard either TFE (14 µM) for ^19^F spectra or TSP (20 µM) for ^1^H spectra, respectively. To assist signal assignment in phosphate buffer conditions 2D ^1^H-^13^C HSQC, ^1^H-^1^H NOESY (800 ms mixing) and ^1^H-^1^H COSY experiments have been recorded at 318 K on a 5 mm TCI cryoprobe.

### 2.4 Production of recombinant mTSPO

mTSPO expression and purification were performed as previously described [19,20]. Briefly, *E. coli* BL21 DE3 strain bacteria containing pET15 plasmid with cDNA of mTSPO were grown in LB medium. mTSPO was extracted from inclusion bodies with SDS 1% and purified by IMAC (ion metal affinity chromatography) with imidazole as competitor. Elution buffer used was 50 mM HEPES, pH 7, 150 mM NaCl and 300 mM imidazole. The protein purity was checked by SDS-PAGE and the protein concentration quantified at 280 nm using the following extinction coefficient 3.88 mL mg^-1^ cm^-1^.

## 3. Results

### 3.1 Synthesis of the fluorinated compound

Bourguignon’s group has identified a L-phenylalanine derivative, NCS7083 (Scheme 1), as a high affinity ligand for TSPO (IC50 = 7 nM) [23]. The replacement of a hydrogen atom by fluorine is commonly used in medicinal chemistry to modulate metabolism or off-target activity. Here, we replaced the phenyl alanine amino acid by its derivative 4-fluoro-L-phenyl alanine, allowing to use the fluorine atom as a sensor for NMR analysis. Both NCS7083 and RF3458 were synthesized in 3 steps (Scheme 1).

### 3.2 Characterization of the fluorinated compound

Chemical shifts of any fluorinated compound needs a reference. TFA is usually used as an internal reference in organic solvents but, being an acid, this reference is not preferred when combined with biological samples. Thus, for the NMR experiments in phosphate buffer (40 mM phosphate pH 6.5, 0.2% DPC, 10% D_2_O), we used TFE as an internal reference for the ^19^F and TSP as internal reference for ^1^H (Figure 1A and 1B) spectra. ^19^F spectrum shows two peaks suggesting the presence of 2 conformers at a ratio of 90 and 10%. ^1^H spectrum also shows two N-CH_3_ signals, at similar ratios, as for the ^19^F spectrum. 2D ^1^H-^13^C HSQC, ^1^H-^1^H COSY and ^1^H-^1^H NOESY (800 ms mixing) spectra (Figure 1C-E) have been exploited to assign the different ^1^H and ^13^C resonances of the NCS7083 and RF3458 compounds at 100 µM in 40 mM phosphate buffer pH 6.5, condition which favours the 2 conformers populated at 90 and 10%, respectively. The ^13^C*α* resonance being close to water signal at room temperature, spectra have also been recorded at 318 K to facilitate its assignment. No clear change in the N-CH_3_ population ratio has been observed between 288 and 318 K. Under these low concentration conditions, no 1D ^13^C spectrum could be recorded. Therefore, only the carbons attached to hydrogens could be assigned via the 2D ^1^H-^13^C HSQC (Figure 1C). The ^1^H signals of the phenyl group holding the fluorine-19 were identified by comparing ^1^H spectra with and without ^19^F-decoupling.

**Figure 1:**
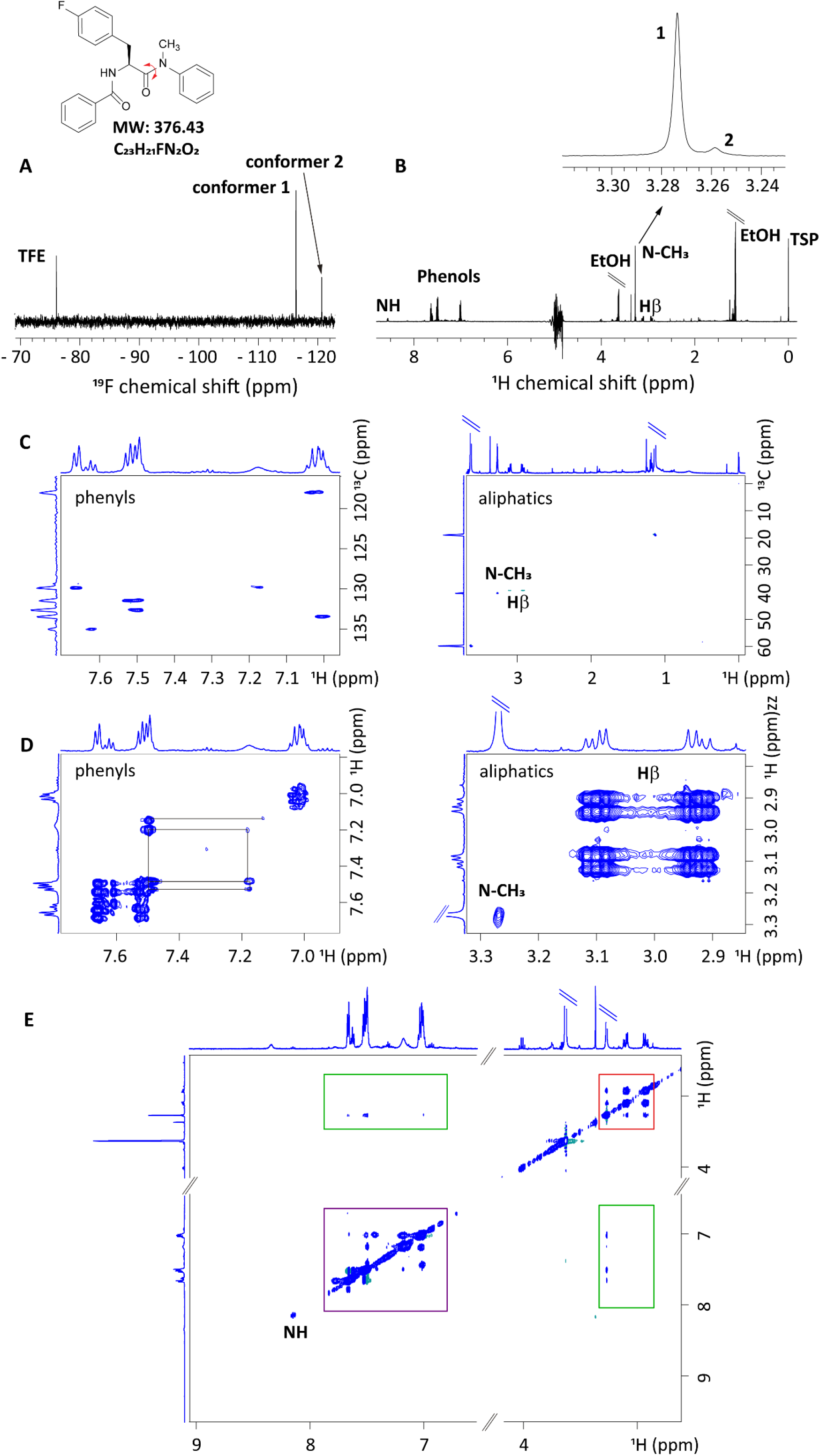
NMR 1D and 2D spectra of ^19^F-RF3458 (100 µM) in phosphate buffer with TFE (15 µM) and TSP (20 µM). (A) 1D ^19^F spectrum recorded at 298 K, showing the TFE peak (−76.6 ppm) and the 2 peaks corresponding to the two conformers of ^19^F-RF3458 (−116.5 and −119.8 ppm). (B) Full ^1^H spectrum recorded at 298 K, shows the various regions (NH, phenyls, N-CH_3_ and H*β*) as well as solvent signals (EtOH and H_2_O) and internal reference TSP; inset highlights the two peaks for the N-CH_3_ group corresponding to the two conformers. (C, D and E) 2D HSQC ^13^C-^1^H, COSY ^1^H-^1^H NOESY ^1^H-^1^H spectra recorded at 318 K. (C), left and right panels correspond to phenyl and aliphatic (N-CH_3_ and H*β*) regions, respectively. (D) COSY ^1^H-^1^H spectrum show cross correlations in-between phenyls (left panel) and H*β* (right panel). (E) NOESY ^1^H-^1^H highlights the cross correlations involving phenyl and aliphatic regions. Green boxes correspond to through space contacts between phenyls and N-CH_3_. Diagonal boxes correspond to through space contacts between N-CH_3_ and H*β* (red box) and between phenyls (purple box).

In order to manage protein-ligand interactions by ^19^F NMR, we have to be in the accessible protein concentrations range (at most in the sub millimolar concentration). Thus, we assessed the ligand concentrations available in our experimental conditions and observed that 50 µM are easily accessible in ethanol (Figure 2A). It has to be noticed that the chemical shift of the ligand is not depending of the concentration. In organic solvents (EtOH or CDCl_3_), only one ^19^F peak was observed (Figure 2A and S2C), whereas in the aqueous media (Figure 2B) used to stabilize the mTSPO [21,22], two peaks are observed suggesting at least two conformations. Moreover, taking into account the referencing compared to the TFE, we noticed that the membrane protein environment (different detergent or detergent-lipid mixture) only affects one of the conformers (Figure 2B). mTSPO protein-fluorinated compound interactions

**Figure 2:**
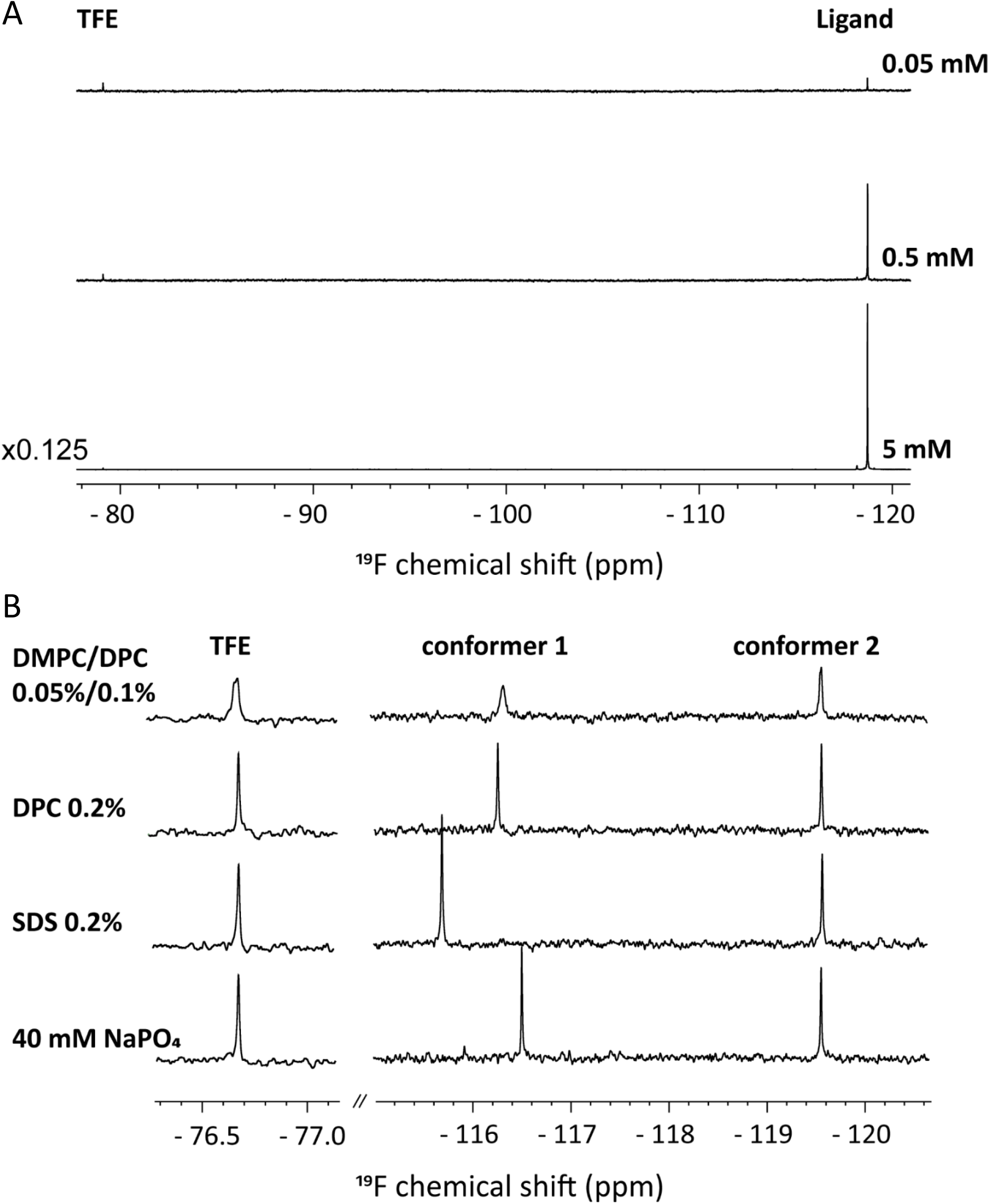
NMR 1D ^19^F spectra of ^19^F-RF3458 in various solvent conditions. (A) Concentration dependence in EtOH (5, 0.5 and 0.05 mM, respectively). (B) Solvent dependence of 100 µM RF3458 in various media (phosphate buffer alone or supplemented with 0.2% SDS or 0.2% DPC or a mixture of DMPC and DPC 0.4/0.2%, respectively). The spectra have been calibrated using TFE resonance as a reference (left panel).

Addition of mTSPO purified in 0.2% SDS to a 100 µM RF3458 ligand solution containing 40 mM phosphate pH 6.5 and 0.2% SDS (Figure 3A) induces a concentration depend chemical shift which follows the same behaviour as the elution buffer used as a control (Figure 3B). It has been previously described that the environment of the mTSPO can be changed from SDS to DPC inducing conformational changes of the protein [19,21]. In 1D ^1^H spectra, we observed the same effect not only for the indole tryptophan region but also for some methyl sidechains (Figure 4A). Thus, we decided to characterize the mTSPO-ligand interactions in the presence of DPC at two concentrations corresponding to half and full-conformational changes (arrows in Figures 4B and 4C).

**Figure 3:**
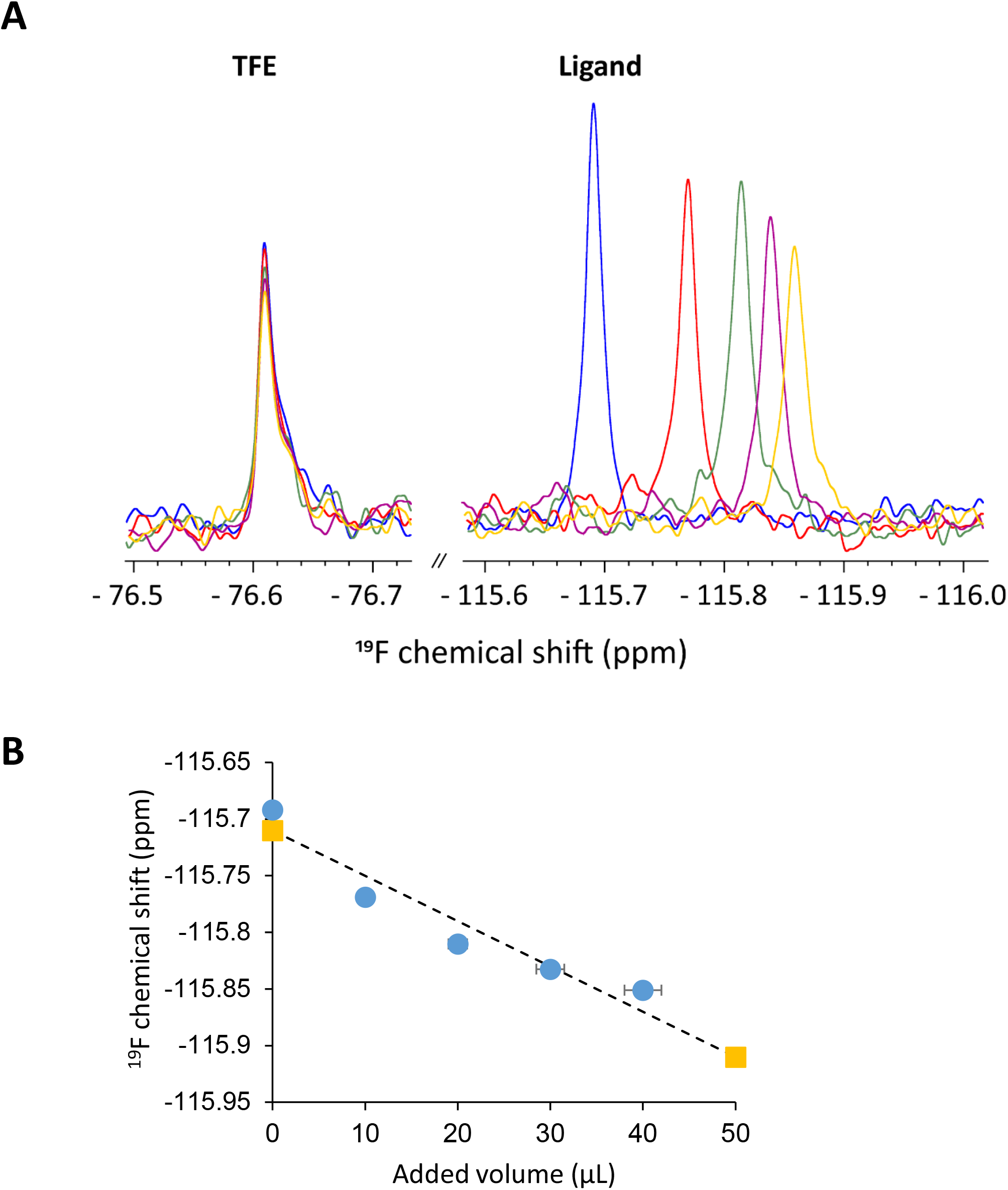
mTSPO titration with ^19^F-RF3458 in 0.2% SDS. (A) NMR 1D ^19^F spectra of 100 µM ^19^F-RF3458 in the presence of increasing concentration of mTSPO purified in 0.2% SDS calibrated using TFE resonance as a reference (left panel). Blue peak corresponds to the free ligand, whereas red, green, purple and yellow peaks correspond to the addition of 10, 20, 30 and 40 µL of mTSPO (∼720 µM) purified in 0.2% SDS, respectively. (B) ^19^F chemical shift evolution as a function of added volumes of mTSPO (grey) or elution buffer (orange).

**Figure 4:**
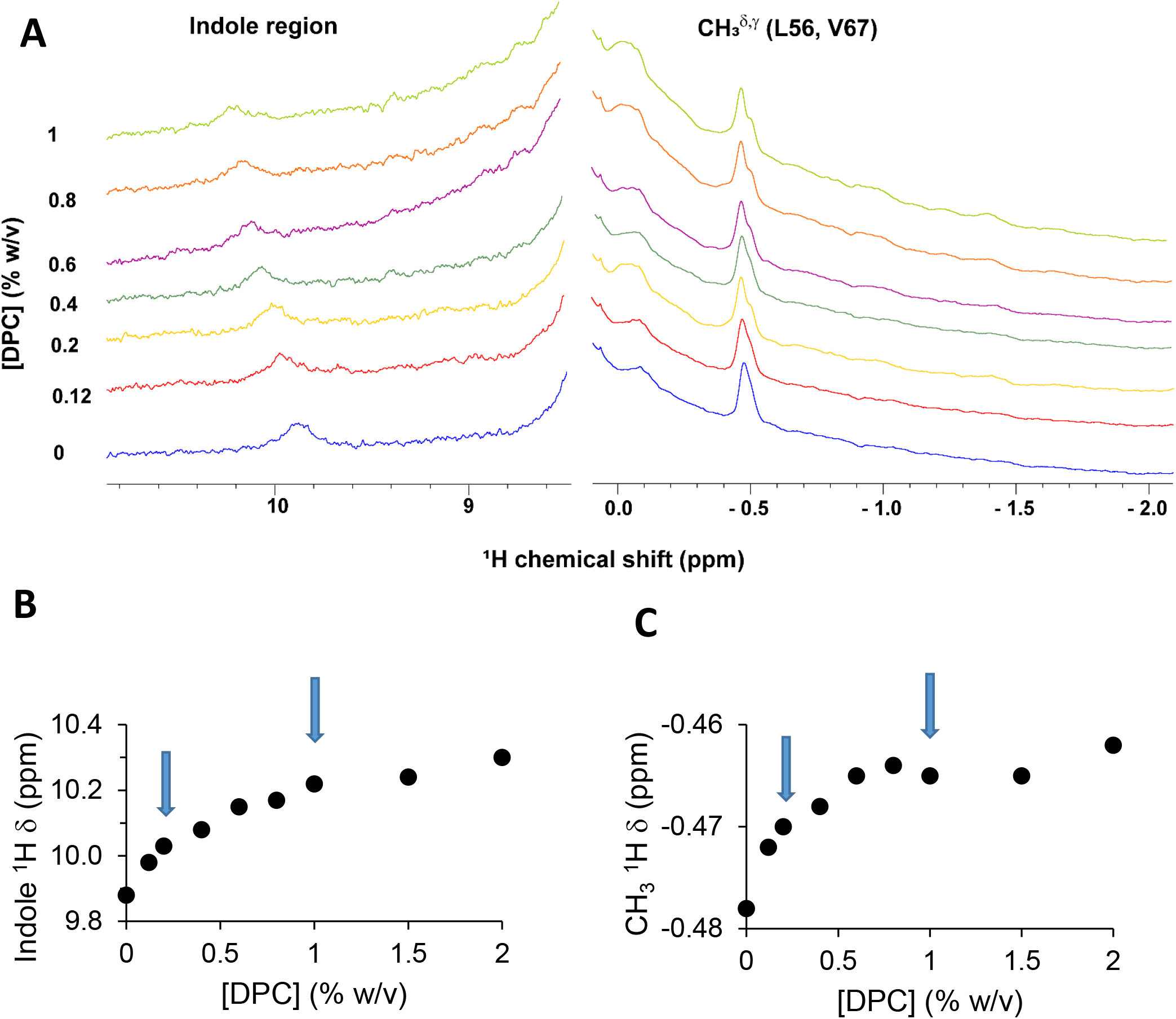
mTSPO titration with DPC. (A) NMR 1D ^1^H spectra of mTSPO (∼40 µM) purified in SDS 0.2% in the presence of increasing amounts of DPC (20 mM phosphate buffer at pH 6.5). ^1^H chemical shift evolution as a function of DPC concentration for indole (B) and methyl (C) regions, respectively. Arrows correspond to the two DPC concentrations used for the titration experiments.

In the presence of DPC 0.2%, addition of mTSPO purified in 0.2% SDS to 50 µM RF3458 induces a concentration depend chemical shift (Figure 5A) which does not follow the trend of the elution buffer used as a control (Figure 5B). The difference between the two experiments (protein versus elution buffer) enabled us to estimate a binding constant of 5 µM (Figure 5C). During the titration experiment, a single ^19^F resonance is observed (central panel), at all the protein-ligand ratios, indicating a rapid exchange between the free and bound forms. The second conformer (right panel) behaves like the first one but the addition of mTSPO induces a smaller chemical shift.

**Figure 5:**
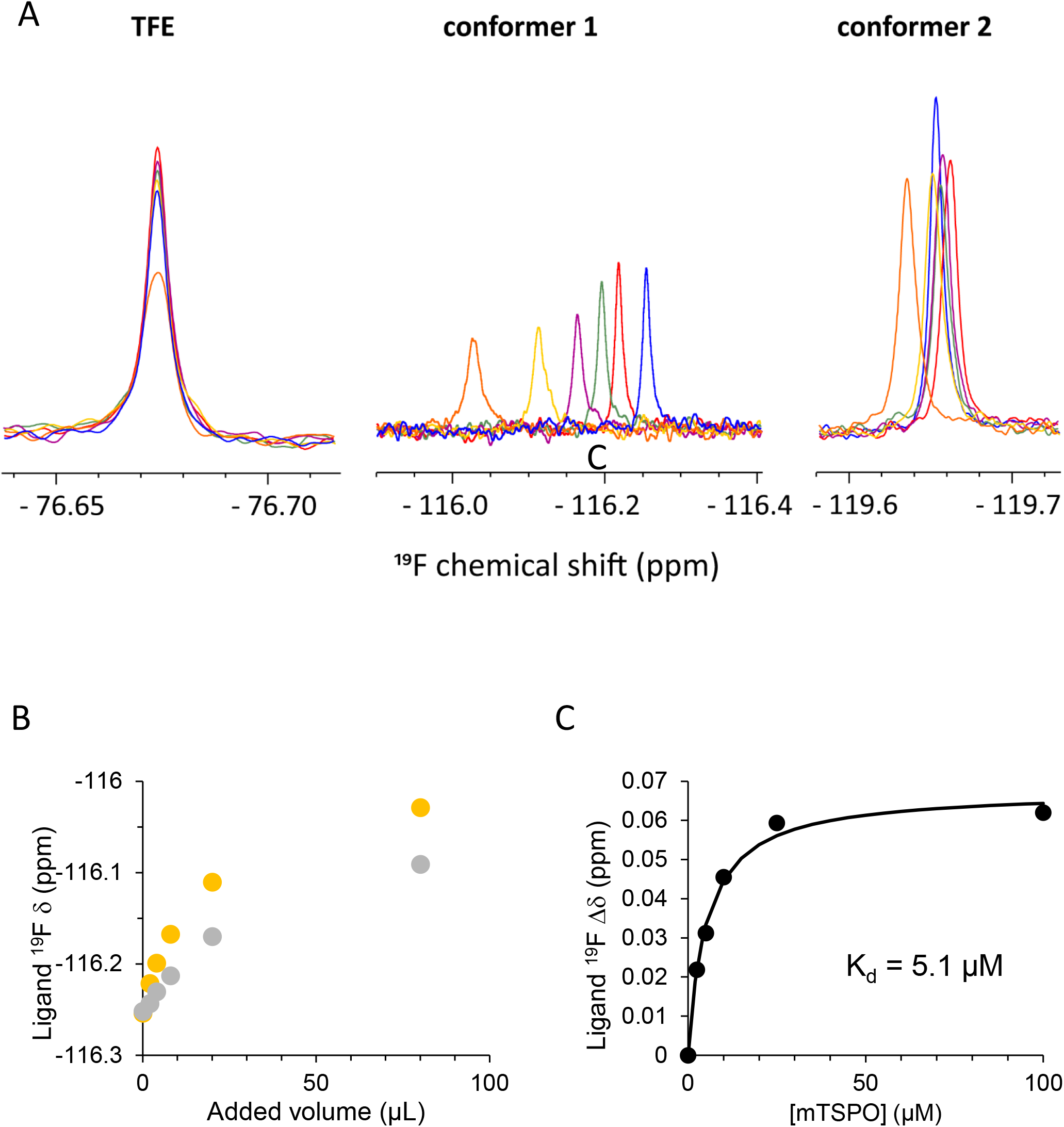
mTSPO titration with ^19^F-RF 3458 in 0.2% DPC. (A) NMR 1D ^19^F spectra of the two conformers of RF3458 at 50 µM in the presence of increasing concentrations of mTSPO purified in 0.2% SDS. Blue resonances correspond to the free forms. (B) ^19^F chemical shift evolution as a function of added volumes of mTSPO (∼650 µM) (grey) or of elution buffer (orange). (C) Binding curve of ^19^F chemical shift difference of the conformer one as a function of mTSPO concentration.

In order to characterize the binding site, we performed replacement experiments with the high affinity drug ligand PK 11195 and we used the (*R*) enantiomer that has the highest affinity [24]. Addition of increasing amounts of (*R*)-PK 11195 shifts the ^19^F-RF3458 signal toward its free form (Figure 6A). We estimate an apparent affinity around 1 mM for (*R*)-PK 11195 in the presence of 50 µM of ^19^F-RF3458 (Figure 6B). This enables to estimate a K_i_, inhibition constant, of ∼90 µM, taking into account the 10-fold excess of ^19^F-RF3458 over its K_d_ and the following equation K_app_ = K_i_*(1+[RF3458]/K_d_). This affinity for PK 11195 is in the range of that measured recently in the same environment by microscale thermophoresis (MST) [22]. The second conformer (right panel) does not behave like the first one since the PK 11195 induces a smaller shift in the opposite direction probably because of the addition of the solvent (DMSO).

**Figure 6:**
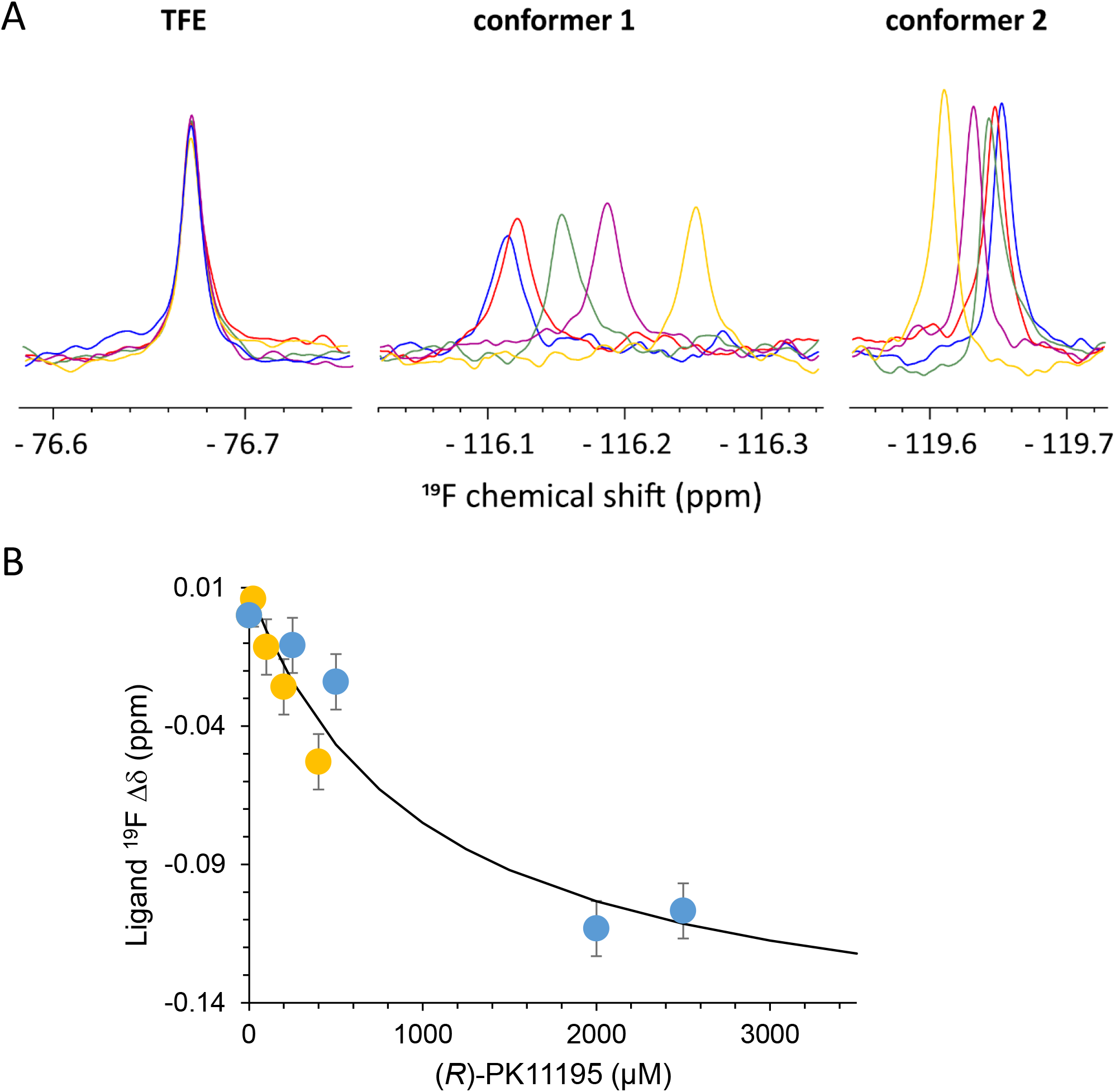
Competition between (*R*)-PK 11195 and ^19^F-RF3458. (A) NMR 1D ^19^F spectra of the first conformer (50 µM bound to 100 µM mTSPO) in the presence of increasing concentration of (*R*)-PK 11195. Blue resonances correspond to the free forms. (B) Binding curve of ^19^F chemical shift difference of the conformer one as a function of (*R*)-PK 11195.

The presence of 1% DPC in the medium changes the ratio between the two conformers (Figure 7A). The interaction with the mTSPO is similar (Figure 7B) to that previously observed at 0.2% DPC. However, the difference between the chemical shift of the ligand in the presence of the protein and of the elution buffer used as a control (Figure 7C), indicates that 1% DPC slightly changes the affinity of the first conformer for the protein, leading to a lower binding constant of ∼25 µM (Figure 7D). A possible explanation is that the ligand is trapped into the detergent micelle making it less accessible for the protein. The second conformer that has a higher intensity compared to 0.2% DPC, still shows a smaller ^19^F chemical shift difference in the mTSPO titration.

**Figure 7:**
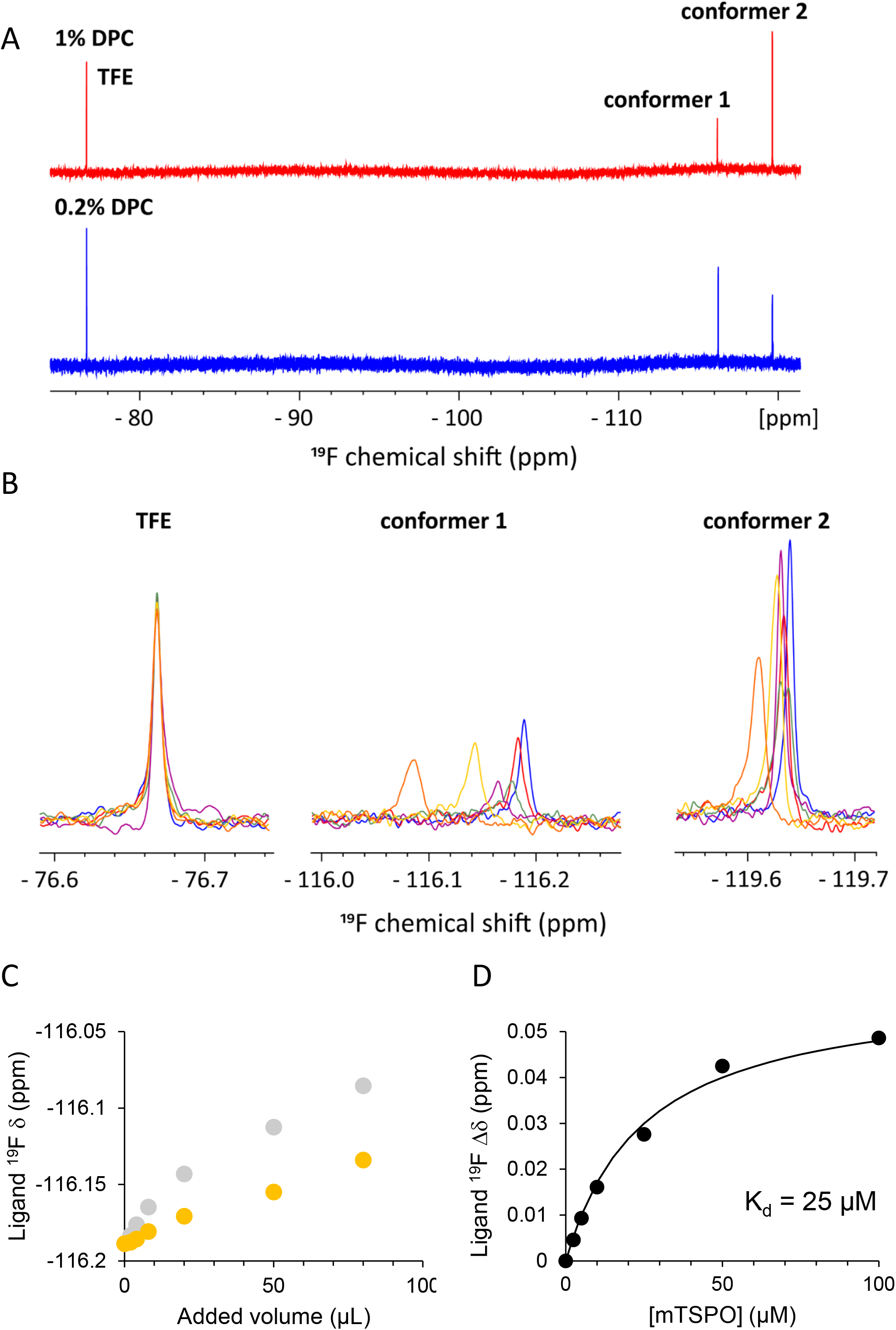
mTSPO titration with ^19^F-RF3458 in 1% DPC. (A) NMR 1D ^19^F spectra of 50 µM RF3458 in the presence 0.2% and 1% DPC. (B) NMR 1D ^19^F spectra of the two conformers in the presence of increasing concentration of mTSPO. Blue resonances correspond to the free forms. (C) ^19^F chemical shift evolution as a function of added volumes of mTSPO (∼650 µM) (grey) or of elution buffer (orange). (D) Binding curve of ^19^F chemical shift difference of the conformer one as a function of mTSPO concentration.

It has been described that the environment of the mTSPO can be changed from SDS to a more native one including lipids, i.e. a DMPC/DPC mixture [22]. It induces conformational changes of the protein affecting secondary and tertiary structure of the mTSPO, that is reflected not only for the indole tryptophan region but also for some methyl sidechains (Figure 8A). Thus, we decided to characterize the mTSPO-ligand interactions in the presence of DMPC/DPC at the concentration corresponding to half of the maximum conformational changes (arrow in Figure 8B, C).

**Figure 8:**
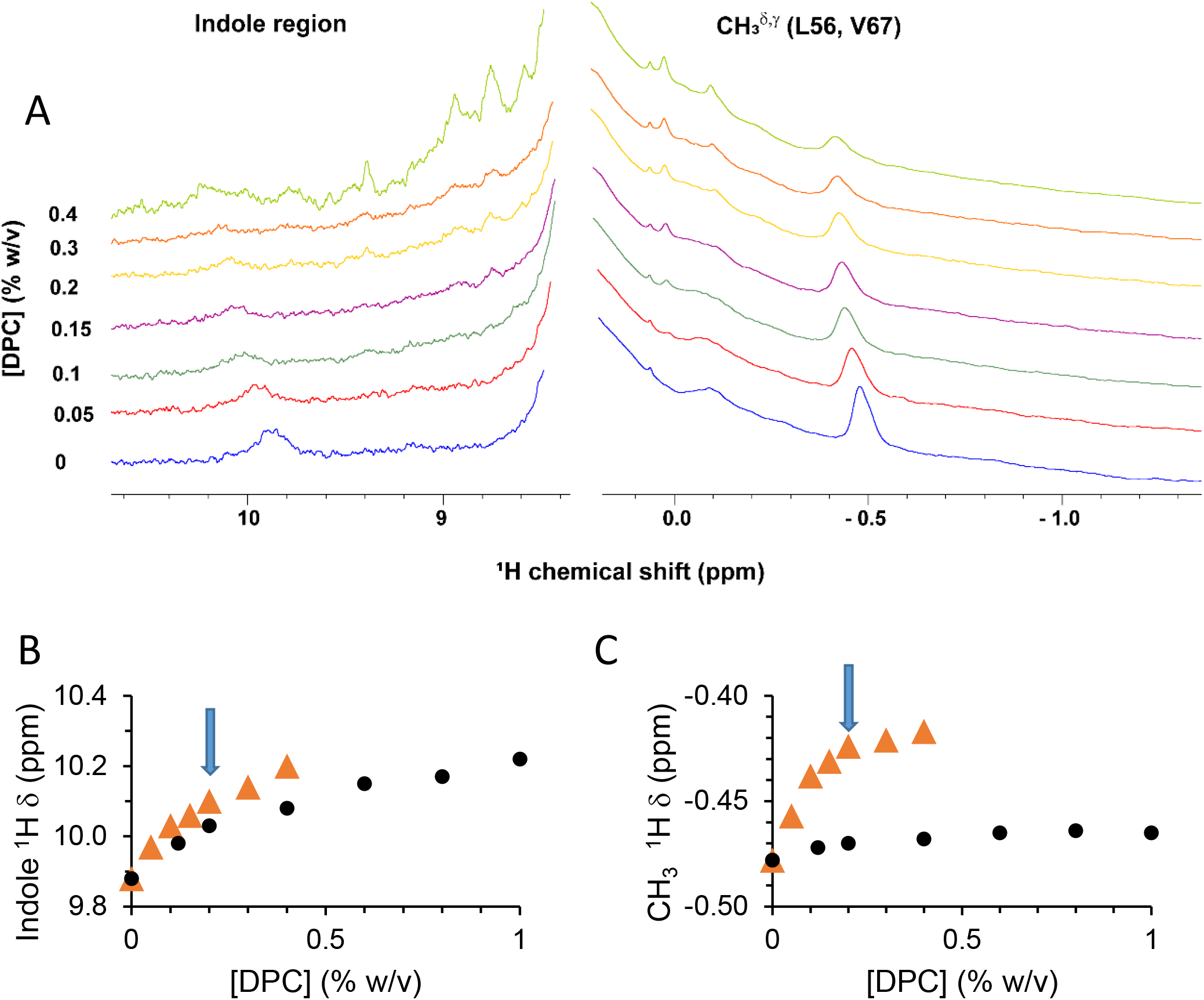
mTSPO titration with a mixture of DMPC/DPC. (A) NMR 1D ^1^H spectra of mTSPO (∼40 µM) purified in SDS 0.2% in the presence of increasing amounts of DMPC/DPC (10%/5%) (20 mM phosphate buffer at pH 6.5). 1H chemical shift evolution as a function of DPC concentration (DMPC/DPC ratio 2/1) for indole (B) and methyl (C) regions, respectively. Arrows correspond to the DMPC/DPC concentration used for the titration experiment.

In the presence of DMPC/DPC 0.4%/0.2% addition of mTSPO purified in 0.2% SDS to 50 µM ligand induces a concentration depend chemical shift (Figure 9A) which does not follow the trend of the elution buffer used as a control (Figure 9B). Contrary to previous measurements in DPC, the titration reveals the migration from one narrow peak of the free form to another narrow peak of the bound form. The intermediate states are characterized by broad resonances which suggest a slow exchange rate between the free and bound states. The difference between the two experiments (protein versus elution buffer) enabled us to estimate a binding constant of 25 µM (Figure 9C). If previous MST studies showed an increase of the affinity for PK 11195 between mTSPO in pure DPC and in DMPC/DPC mixture [22], herein we observed a decrease of the affinity. This might be due to the presence of an excess of DMPC/DPC on top of the SDS together with the mTSPO as previously observed when we increased the DPC from 0.2 to 1%. In other words, the previously reported MST experiments [22] have been conducted with mTSPO purified in DMPC/DPC; hence, with a low concentration of free DMPC/DPC (0.05%/0.1%) and in the absence of SDS, unlike our conditions herein.

**Figure 9:**
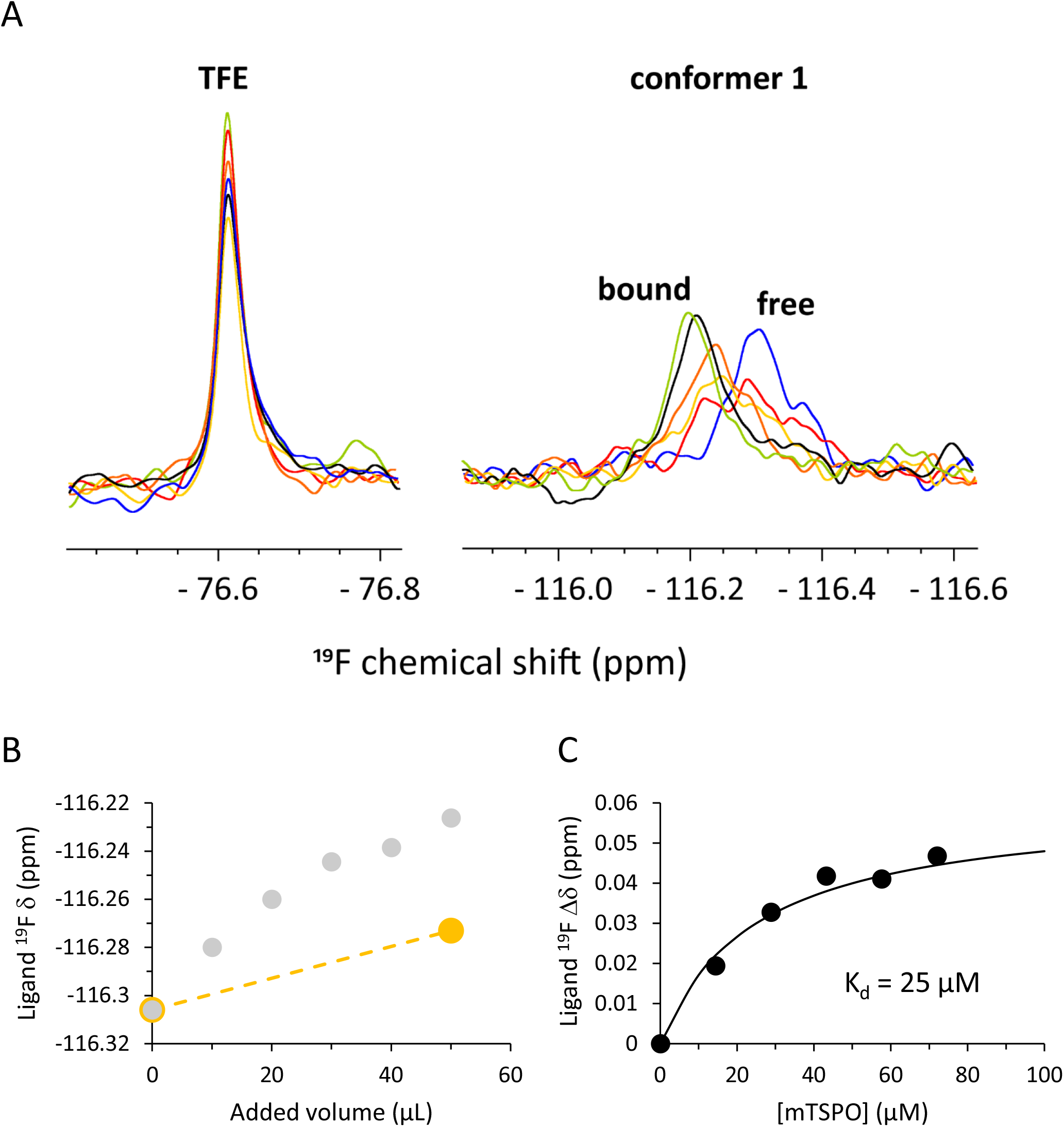
mTSPO titration with ^19^F-RF3458 in a mixture of DMPC/DPC. (A) NMR 1D ^19^F spectra of 100 µM RF3458 in the presence of DMPC/DPC 0.4/0.2% in the presence of increasing concentrations of mTSPO purified in 0.2% SDS. Blue resonances correspond to the free forms. (B) ^19^F chemical shift evolution as a function of added volumes of mTSPO (∼720 µM) (grey) or of elution buffer (orange). (C) Binding curve of ^19^F chemical shift difference of the conformer one as a function of mTSPO concentration.

## 4. Discussion

Among the endogenous ligands of TSPO, a polypeptide of 9 kDa has been shown to have multiple biological actions [25]. Smaller dipeptides have been designed and demonstrated anxiolytic effects [4,5,7,26]. Based on docking studies, these dipeptides seem to bind the same TSPO cavity identified for the high-affinity drug ligand PK 11195 [7]. However, no direct binding studies and/or affinity measurements for TSPO are currently available.

The new amino acid derived fluorinated ligand presented in this work has been used to measure its affinity for mTSPO in different detergent-containing environments. mTSPO has been previously described as being incompletely folded in SDS [19,21,27] and thus unable to bind its high-affinity ligand PK 11195 [19] in agreement with our observation of only unspecific binding. When immersed in DPC, mTSPO exhibits superior folding characteristics which may facilitate ligand binding, as confirmed by the acquisition of an atomic-level structure of the complex between the high-affinity ligand (*R*)-PK 11195 and mTSPO by solution NMR [28]. Additionally, our ^19^F NMR titration investigations in DPC revealed a distinct downfield shift (high chemical shift values) of the ^19^F resonance, indicating rapid interconversion between the free and bound states. Essentially, this implies a fast off-rate constant (*k*_*off*_), which indicates a low binding affinity. In contrast, when mTSPO is in a mixed lipid-detergent environment, the titration of the ^19^F ligand presents a completely different profile. We observed a decrease in the intensity of the free-form peak and the emergence of a bound-form peak that intensifies as the protein concentration increases. The coexistence of these two peaks suggests a regime of slow exchange, characterized by a slow *k*_*off*_, which is indicative of a high binding affinity. However, the measured affinity is also dependent of the environment such micelles which can trap the ligand. Moreover, the presence of mixture of detergent (SDS and DPC) and lipids might make more complex the interpretation of the data. Further experiments are needed to measure the mTSPO affinities in pure detergent or lipid detergent mixture as previously reported [22]. Nevertheless, we can infer that the ligand accessibility to the mTSPO binding site and its dynamics are governed by the protein’s environment. We have shown that (*R*)-PK 11195 enantiomer is able to replace the ^19^F-RF3458 ligand, suggesting the binding to the same site. Further studies with ^15^N-labelled mTSPO would permit to acquire 2D ^1^H-^15^N HSQC spectra and therefore identify the amino acids specifically involved in the binding of the two ligands. It would also permit to characterize the effects of the different environments upon the structure of the mTSPO in particular the splitting of indoles and methyl side chains [19].

(*R*) and (*S*)-PK 11195 enantiomers [24,29] exhibit different affinities for TSPO [30] but radioactivity cannot measure directly their individual affinity. Indeed, in rat models [30] the binding affinity of the (*R*) enantiomer has been shown to be significantly greater than that of the racemic mixture. Contrary to radioactivity, ^19^F NMR spectroscopy can detect the coexistence of conformers. In our NMR study, we have identified the presence of two conformers for the phenyl alanine derived ligand ^19^F-RF3458. However, the lower intensity and smaller chemical shift displacement upon mTSPO binding as well as the effect of elution buffer did not permit to clearly estimate an affinity for the second conformer.

## 5. Conclusion and perspectives

Fluorine-18 tracers are promising alternatives to the ^11^C-based radioligands for TSPO PET imaging, but little is known about the molecular level of TSPO-ligand interaction for all these tracers. The fluorinated ligand described herein is a phenyl alanine derived molecule which is close to the dipeptide ligands previously described [4,5,7,26]. It has the advantage of being versatile for probe attachment since the fluorine molecule can be attached on any of the 3 phenyl rings or on all them. In addition, the possibility of adding a CF_3_ group instead of the fluorine alone is particularly interesting for increasing the ^19^F NMR signal. What about the affinity of ligand bearing a CF_3_ group instead of a single F, this substitution may lead to a bigger molecule, hence more steric hindrance? In addition, fluorine-18 tracers appear as promising molecules to design new TSPO ligands by screening chemo-library of fragments capable to displace them and to characterize binding site domains.

Transition toward clinical applications, such as PET imaging, would require toxicology studies, determination of possible cellular targets, and cellular accessibility for this hydrophobic ligand that has three phenyls rings, in particular if the brain is targeted since it has to cross the hematoencephalic barrier.

## Supporting information

SSupplemental

## Author contributions

LD, FB and JJL: Conceptualization, Data curation, Formal analysis, Methodology, Validation, Writing – review & editing; SS and CH: Methodology; AM: Methodology, Formal analysis, Validation.

## Acknowledgments

We would like to thank all our colleagues for the exciting discussions during the monthly meetings of the French-speaking TSPO group that was initiated in 2017. Financial support from CNRS (LD, AM, FB, JJL), Université de Reims Champagne-Ardenne (LD, AM, CH), Université de Strasbourg (SS, FB) and Sorbonne Université (JJL) is acknowledged. We acknowledge Mohyeddine Taleb for technical help in the expression and the purification of mTSPO at the beginning of the project and Claire Troufflard for technical support on the Jussieu NMR platform. Stimulating discussions about ^19^F NMR experiments with Emeric Miclet and Ewen Lescop were highly appreciated.

## Abbreviations

BzCl: Benzoyl chloride
COSY: correlation spectroscopy
DBI: diazepam-binding inhibitor
DBU: 1,8-diazabicyclo 5.4.0 undec-7-ene
DCM: Dichloromethane
DIEA: Diisopropylethylamine
DMF: Dimethylformamide
DMPC: 1,2-dimyristoyl-sn-glycero-3-phosphocholine
DPC: dodecylphosphocholine
EtOAc: ethyl acetate
HATU: hexafluorophosphate azabenzotriazole tetramethyl uronium
HEPES: 2-[4-(2-hydroxyethyl)piperazin-1-yl]ethanesulfonic acid
HPLC: high-performance liquid chromatography
HSQC: Heteronuclear Single Quantum Coherence
IMAC: immobilized metal affinity
LB: Luria-Bertani broth
MST: microscale thermophoresis
MRI: magnetic resonance imagining
NOESY: nuclear Overhauser effect spectroscopy
NMR: nuclear magnetic resonance
PET: positron electron tomography
PK 11195: 1-(2-Chlorophenyl)-N-methyl-N-(1-methylpropyl)-3-isoquinolinecarboxamide
PK 14105: N-Butan-2-yl-1-(2-fluoro-5-nitrophenyl)-N-methylisoquinoline-3-carboxamide
SDS: sodium dodecyl sulfate
TFA: trifluoroacetic acid
TFE: 2,2,2-trifluoroethanol
TSP: sodium trimethylsilyl propionate
TSPO: translocator protein

