## Supplementary material for "Fluor NMR study of amino acid derived ligand to study TSPO": SSupplemental

**Acronyms**

DAD: diode-array detection

HR-MS: high-resolution mass spectrometry

NMR: nuclear magnetic resonance

RP-HPLC-MS: reverse phase high-performance liquid chromatography-mass spectrometry

TLC: thin layer chromatography

**Materials and Instruments**

All commercial reagents were used without purification. Analytical TLC was performed using Merck 60 F254 silica gel plates, and visualized by exposure to ultraviolet light (254 nm). Compounds were purified on silica gel Merck 60 (particle size 0.040−0.063 nm). NMR spectra were recorded on Bruker Avance III or Neo spectrometers operating at 600, 500 or 400 MHz for ^1^H, 150, 126 or 101 MHz for ^13^C, and 470 MHz for ^19^F. All chemical shift values δ and coupling constants *J* are quoted in ppm and in Hz, respectively, and multiplicity are quoted using: s = singlet, d = doublet, t = triplet, q = quartet, m = multiplet, and br = broad. Analytical RP-HPLC-MS was performed using a LC 1200 Agilent with quadrupole-time-of-flight (QTOF) (Agilent Accurate Mass QToF 6520) with a ZORBAX Agilent C18-column (C18, 50 mm × 2.1 mm; 1.8 μm) using the following parameters: (1) the solvent system: A (0.05% of formic acid in acetonitrile) and B (0.05% of formic acid in H_2_O); (2) a linear gradient: t = 0 min, 98% B; t = 8 min, 0% B; t = 12.5 min, 0% B; t = 12.6 min, 98% B; t = 13 min, 98% B; (3) flow rate of 0.5 mL/min; (4) column temperature: 35 °C; (5) DAD scan from 190 to 700 nm; and (6) ionization mode: ESI+. HPLC were performed using a Dionex UltiMate 3000 using the following parameters: column WATERS XSelect CSH C18 (5µm, 4.6 x 150 mm); temperature: 40°C; flow rate = 2 mL/min; eluent system: A (0.05% of TFA in H_2_O) and B (MeCN) with 5 to 100% of B in 10 min.

NCS7083 (**4a**) ^1^H NMR (600 MHz, CDCl_3_) δ_H_ 2.87 (1H, dd, ^3^*J*_Hβ',Hα_ = 6.40 Hz, ^2^*J*_Hβ',Hβ_ = 13.53 Hz), 3.01 (1H, dd, ^3^*J*_Hβ,Hα_ = 7.66 Hz, ^2^*J*_Hβ,Hβ’_ = 13.53 Hz), 3.24 (3H, s), 5.05 (1H, m, ^3^*J*_Hβ',Hα_ = 6.40 Hz, ^3^*J*_Hβ,Hα_ = 7.70 Hz, ^3^*J*_NH,Hα_ = 8.18 Hz), 6.84 (1H, d, *J* = 8.30), 6.98 (2H, m), 6.99 (2H, m), 7.23 (3H, dd, *J* = 3.09/3.22 Hz), 7.35 (3H, m), 7.41 (2H, t, *J* = 7.35 Hz), 7.48 (1H, t, *J* = 7.85 Hz), 7.74 (2H, d, *J* = 8.30 Hz) (Figure S1A); ^13^C NMR (150 MHz, CDCl_3_) δ_C_ 37.8 (s, N-CH_3_), 39.5 (s, Cβ), 51.6 (s, Cα), 127.0 (s, Cc green), 127.2 (s, Ca,a’ magenta), 128.3 (s, Cc blue), 128.6 (s, Cb,b’ magenta), 128.9 (s, Cb,b’ green), 129.6 (s, Ca,a’ green), 129.9 (s, Ca,a’ blue), 131.7 (s, Cc magenta), 134.2 (s, Cd magenta), 136.4 (s, Cd green), 142.5 (s, Cd blue), 166.6 (s, C=O-NH), 171.7 (s, C=O-NH_3_) (Figure S1B); ^15^N NMR (120 MHz, CDCl_3_) δ_N_ 114.8 (NH), 125.6 (N-CH_3_). ^1^H & ^13^C NMR of NCS7083 (**4a**) were in agreement with the one previously reported (C. Houyvet, PhD thesis).

RF3458 (**4b**) ^1^H NMR (600 MHz, CDCl_3_) δ_H_ 2.83 (1H, dd, ^3^*J*_Hβ',Hα_ = 6.31 Hz, ^2^*J*_Hβ',Hβ_ = 13.73 Hz), 2.98 (1H, dd, ^3^*J*_Hβ,Hα_ = 7.55 Hz, ^2^*J*_Hβ,Hβ’_ = 13.73 Hz), 3.25 (3H, s), 5.06 (1H, m, ^3^*J*_Hβ',Hα_ = 6.31 Hz, ^3^*J*_Hβ,Hα_ = 7.55 Hz, ^3^*J*_NH,Hα_ = 8.04 Hz), 6.84 (1H, d, *J* = 8.14), 6.93 (6H, m), 6.94 (6H, m), 7.37 (3H, m), 7.39 (3H, d, *J* = 8.12), 7.42 (2H, t, *J* = 7.16 Hz), 7.49 (2H, t, *J* = 7.61 Hz), 7.74 (2H, d, *J* = 7.69 Hz) (Figure S2A); ^13^C NMR (150 MHz, CDCl_3_) δ_C_ 37.8 (s, N-CH_3_), 38.7 (s, Cβ), 51.5 (s, Cα), 115.4 (d, Cb,b’ green, ^2^*J*_CF_ = 21.27 Hz), 127.2 (s, Ca,a’ magenta), 127.4 (s, Cb,b’ blue), 128.7 (s, Cc magenta), 130.0 (s, Ca,a’ blue), 131.0 (d, Ca,a’ green, ^3^*J*_CF_ = 7.98 Hz), 131.8 (s, Cb,b’ magenta), 132.2 (d, Cd green, ^4^*J*_CF_ = 3.17 Hz), 134.1 (s, Cd magenta), 142.4 (s, Cd blue), 134.2 (s, Cd magenta), 136.4 (s, Cd green), 142.5 (s, Cd blue), 162.1 (d, Cc green, ^1^*J*_CF_ = 245.11 Hz), 166.5 (s, C=O-NH), 171.6 (s, C=O-NH_3_) (Figure S2B); ^15^N NMR (120 MHz, CDCl_3_) δ_N_ 114.8 (NH), 125.6 (N-CH_3_). ^19^F NMR (470 MHz, CDCl_3_) δ -115.95 (Figure S2C). HR-MS (ESI+): m/z calculated for C_23_H_21_FN_2_O_2_ [M+H]^+^ 377.1665, found 377.1683.

**Table S1**: ^1^H, ^13^C, ^15^N and ^19^F chemical shifts (in ppm), integrals and scalar couplings of NCS7083 and RF3458 in CDCl_3_.

|  | NCS7083 (C_23_H_22_N_2_O_2_) | | | RF3458 (C_23_H_21_FN_2_O_2_) | | |
| --- | --- | --- | --- | --- | --- | --- |
| Name | δ_H_ (ppm) | nH, type, *J* (Hz) | δ_C/N_ (ppm) | δ_H/F_ (ppm) | nH, type, *J* (Hz) | δ_C/N_ (ppm) |
| CαHα | 5.05 | 1H, m, 6.40, 7.70, 8.18 | 51.6 | 5.06 | 1H, m, 6.31, 7.55, 8.04 | 51.5 |
| CβHβ | 3.01 | 1H, dd, 7.70, 13.53 | 39.5 | 2.98 | 1H, dd, 7.55, 13.73 | 38.7 |
| CβHβ’ | 2.87 | 1H, dd, 6.40, 13.53 | 39.5 | 2.83 | 1H, dd, 6.31, 13.73 | 38.7 |
| NH | 6.84 | 1H, d, 8.30 | -/114.8 | 6.84 | 1H, d, 8.14 | -/114.0 |
| N-CH_3_ | 3.24 | 3H, s | 37.8/125.6 | 3.25 | 3H, s | 37.8/125.6 |
| C=O-NH | - | - | 166.6 | - | - | 166.5 |
| C=O-N-CH_3_ | - | - | 171.7 | - | - | 171.6 |
| Ca,a’Ha,a’ | 7.74 | 2H, d, 8.3 | 127.2 | 7.74 | 2H, d, 7.69 | 127.2 |
| Cb,b’Hb,b’ | 7.41 | 2H, t, 7.35 | 128.6 | 7.42 | 2H, t, 7.16 | 131.8 |
| CcHc | 7.48 | 2H, t, 7.85 | 131.7 | 7.49 | 2H, t, 7.61 | 128.7 |
| Cd | - | - | 134.2 | - | - | 134.1 |
| Ca,a’Ha,a’ | 7.35 | 3H, m | 129.9 | 7.39 | 3H, d, 8.12 | 130 |
| Cb,b’Hb,b’ | 6.98 | 2H, m | 127.4 | 6.94 | 6H, m | 127.4 |
| CcHc | 7.35 | 3H, m | 128.3 | 7.37 | 3H, m | 128.4 |
| Cd | - | - | 142.5 | - | - | 142.4 |
| Ca,a’Ha,a’ | 6.99 | 2H, m | 129.6 | 6.93 | 6H, m | 131.0, ^3^*J*_CF_ = 7.98 |
| Cb,b’Hb,b’ | 7.23 | 3H, t | 128.6 | 6.93 | 6H, m | 115.4, ^2^*J*_CF_ = 21.27 |
| CcHc | 7.23 | 3H, t | 127.0 | - | - | 162.1, ^1^*J*_CF_ = 245.11 |
| Cd | - | - | 136.4 | - | - | 132.2, ^4^*J*_CF_ = 3.17 |
| F | - | - | - | -115.95 | - | - |

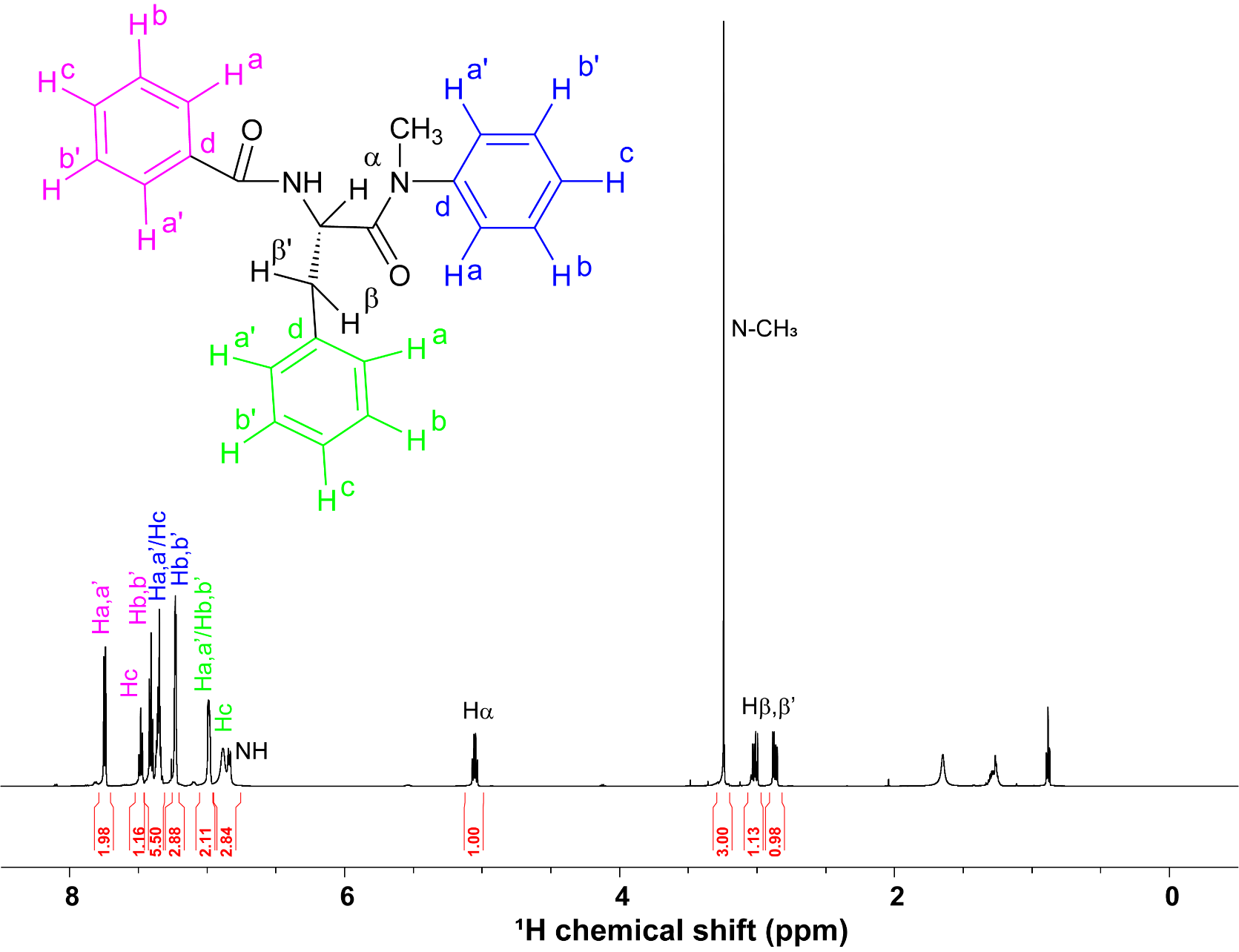

Figure S1A: NMR ^1^H spectrum of NCS7083 (~12 mg/mL) in CDCl_3_ at 298 K. Spectrum recorded on a Bruker Avance III spectrometer (600 MHz ^1^H Larmor frequency) equipped with a 5 mm TCI probe head. Residual protonated solvent signal at 7.26 ppm has been used as an internal reference.

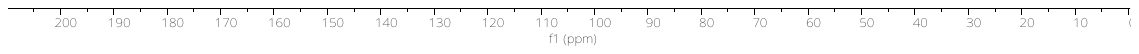

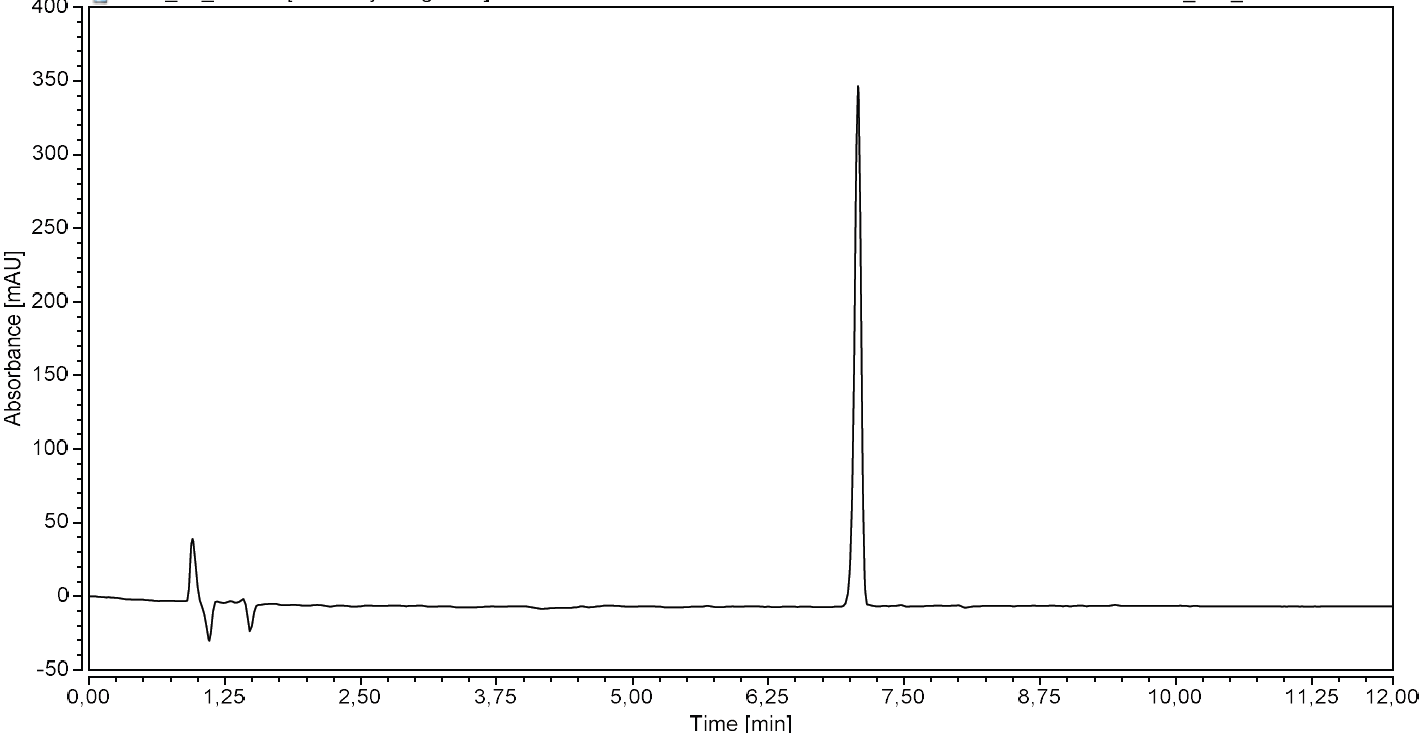

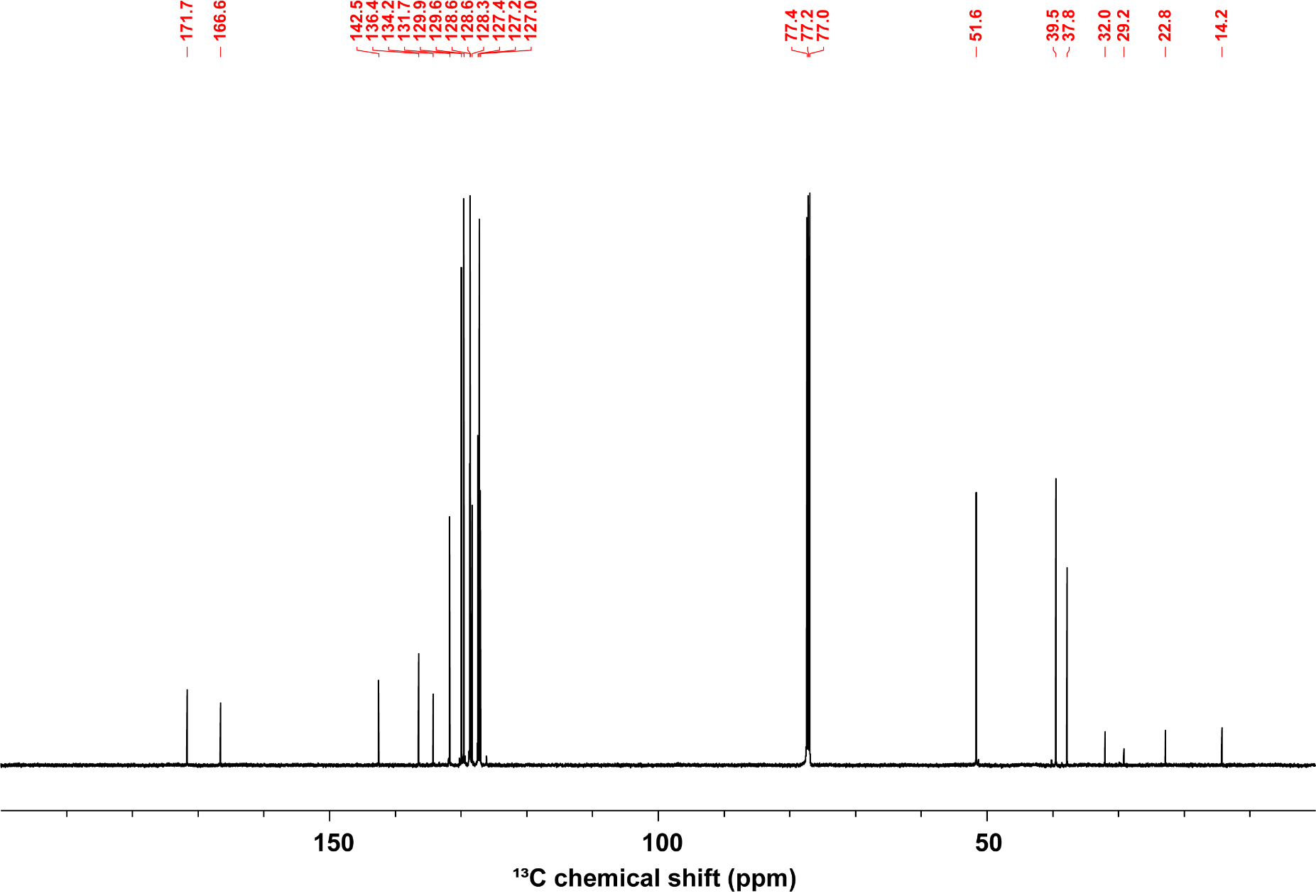

B) Cpd **4a** - ^13^C NMR (126 MHz, CDCl_3_)

C) Cpd **4a** - HPLC Purity (210 nm)

Figure S1B: NMR ^13^C spectrum of NCS7083 (~12 mg/mL) in CDCl_3_ at 298 K. Spectrum recorded on a Bruker Avance III spectrometer (150 MHz ^13^C Larmor frequency) equipped with a 5 mm TCI probehead. Residual solvent signal at 77.16 ppm has been used as an internal reference.

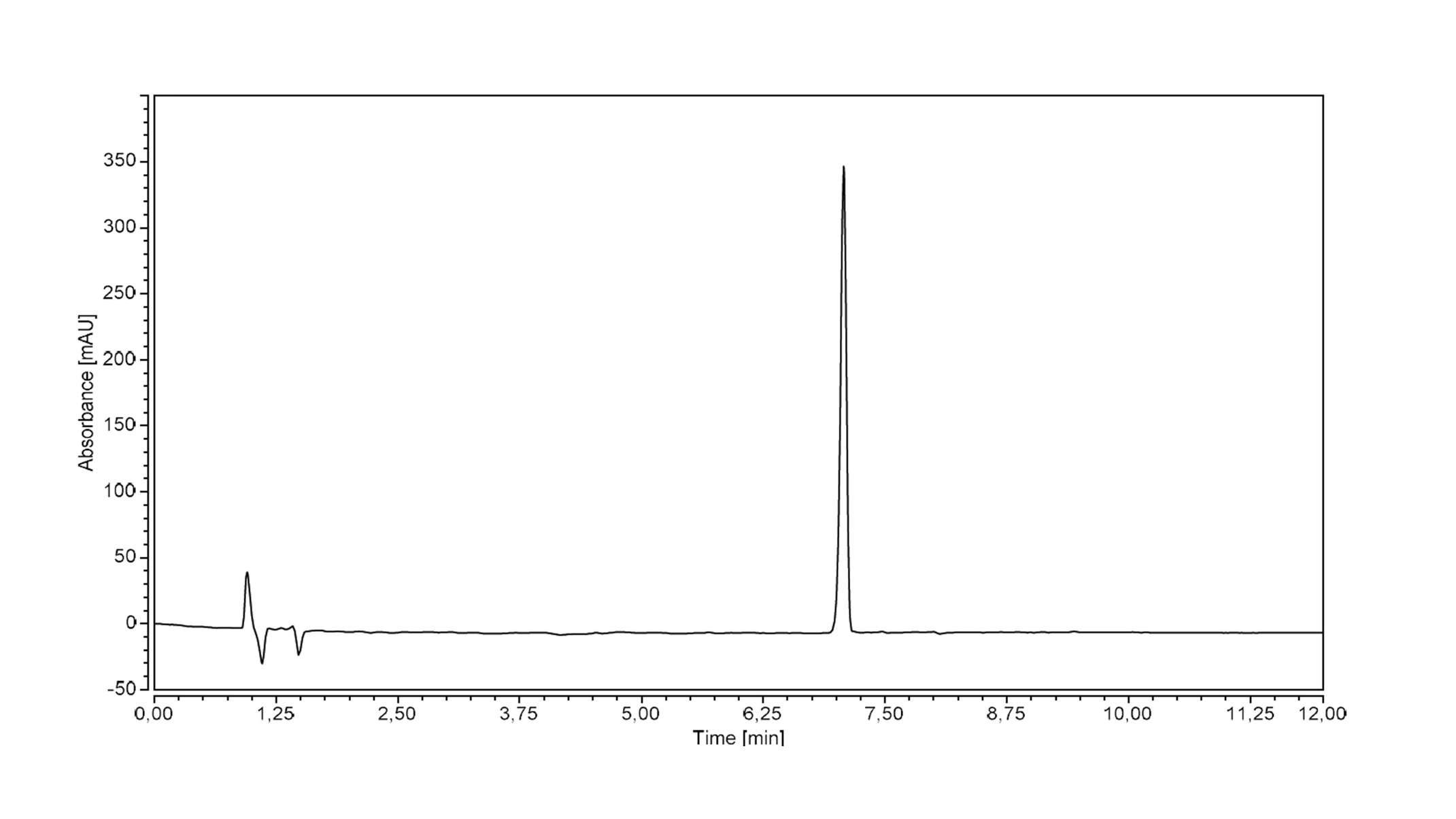

Figure S1C: HPLC purity (210 nm) of NCS7083.

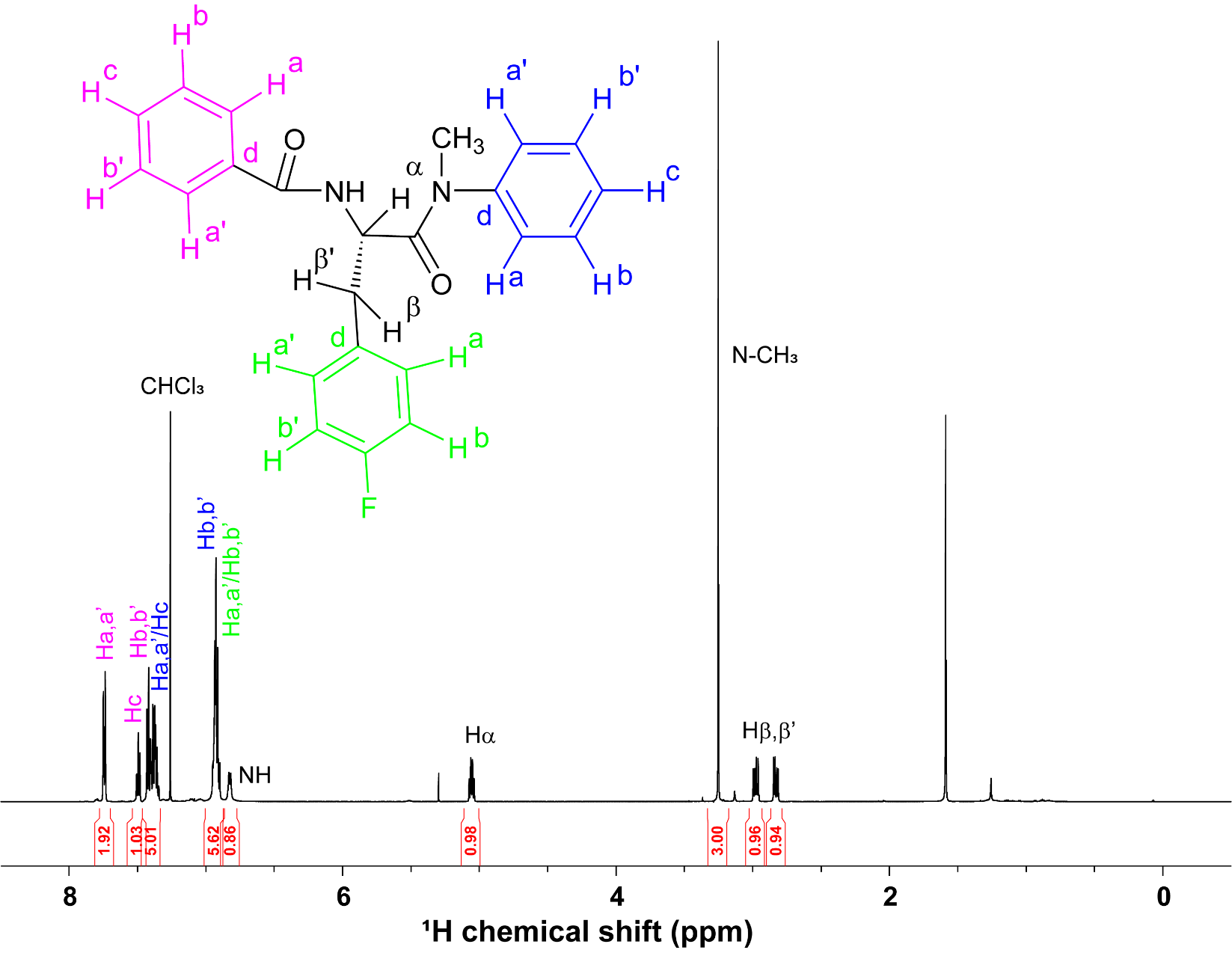

Figure S2A: NMR ^1^H spectrum of RF3458 (~10 mg/mL) in CDCl_3_ at 298 K. Spectrum recorded on a Bruker Avance III spectrometer (600 MHz ^1^H Larmor frequency) equipped with a 5 mm TCI probehead. Residual protonated solvent signal at 7.26 ppm has been used as an internal reference.

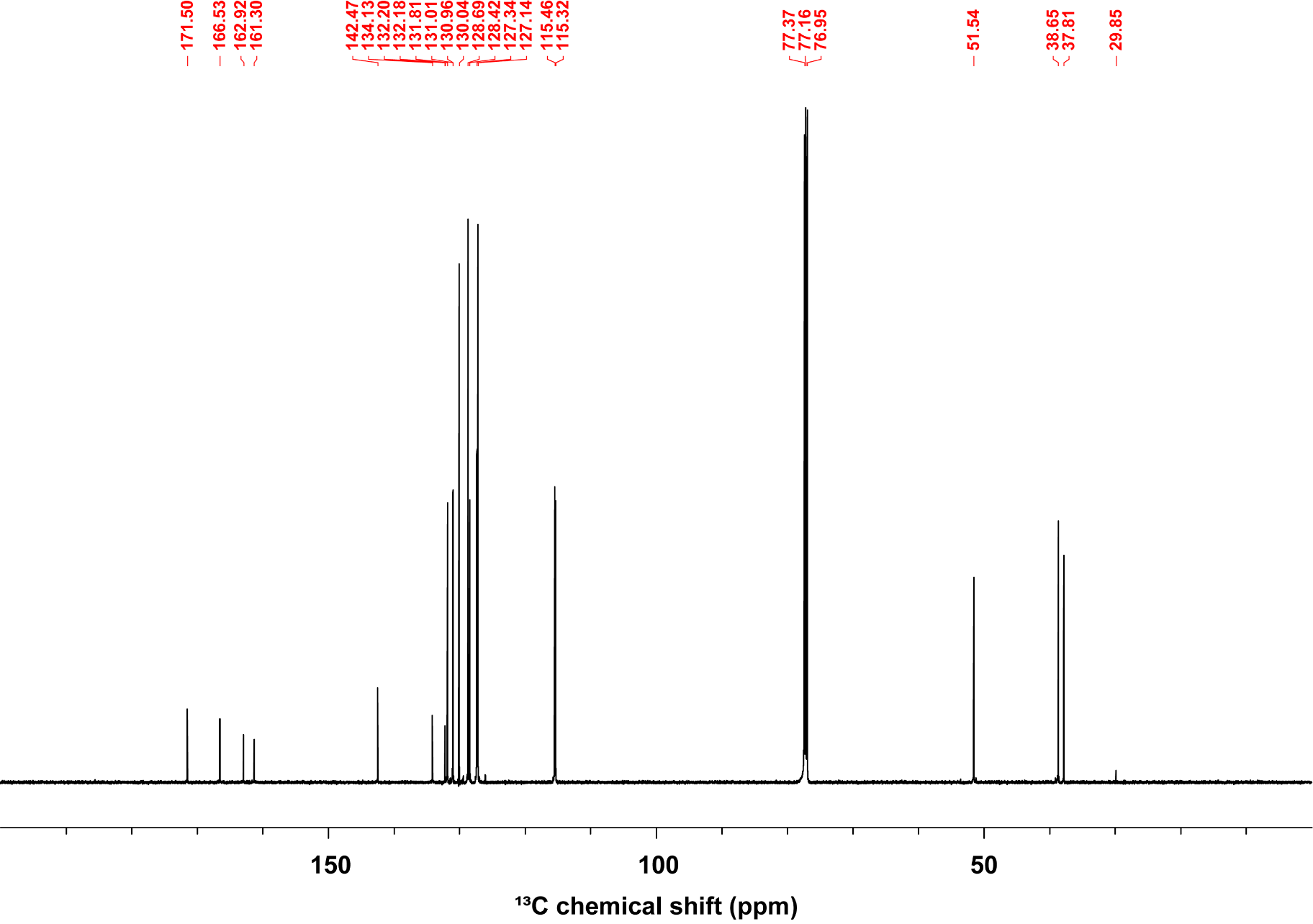

Figure S2B: NMR ^13^C spectrum of RF3458 (~10 mg/mL) in CDCl_3_ at 298 K. Spectrum recorded on a Bruker Avance III spectrometer (150 MHz ^13^C Larmor frequency) equipped with a 5 mm TCI probe head. Residual solvent signal at 77.16 ppm has been used as an internal reference.

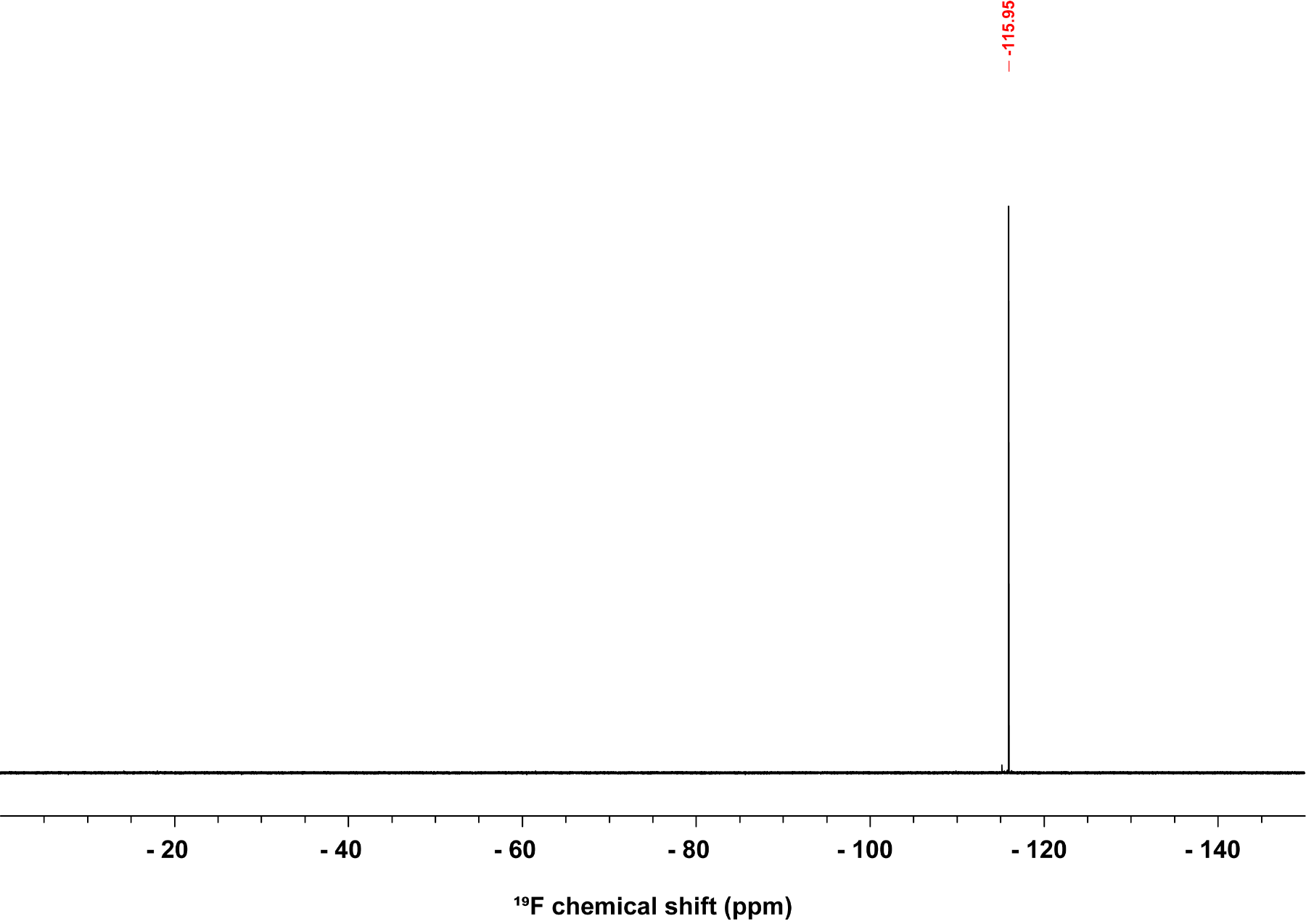

Figure S2C: NMR ^19^F spectrum of RF3458 (~10 mg/mL) in CDCl_3_ at 298 K. Spectrum recorded on a Bruker Neo spectrometer (470 MHz ^19^F Larmor frequency) equipped with a 5 mm iprobe. ^19^F chemical shift is relative to the external reference of α,α,α-trifluorotoluene at -63.72 ppm.

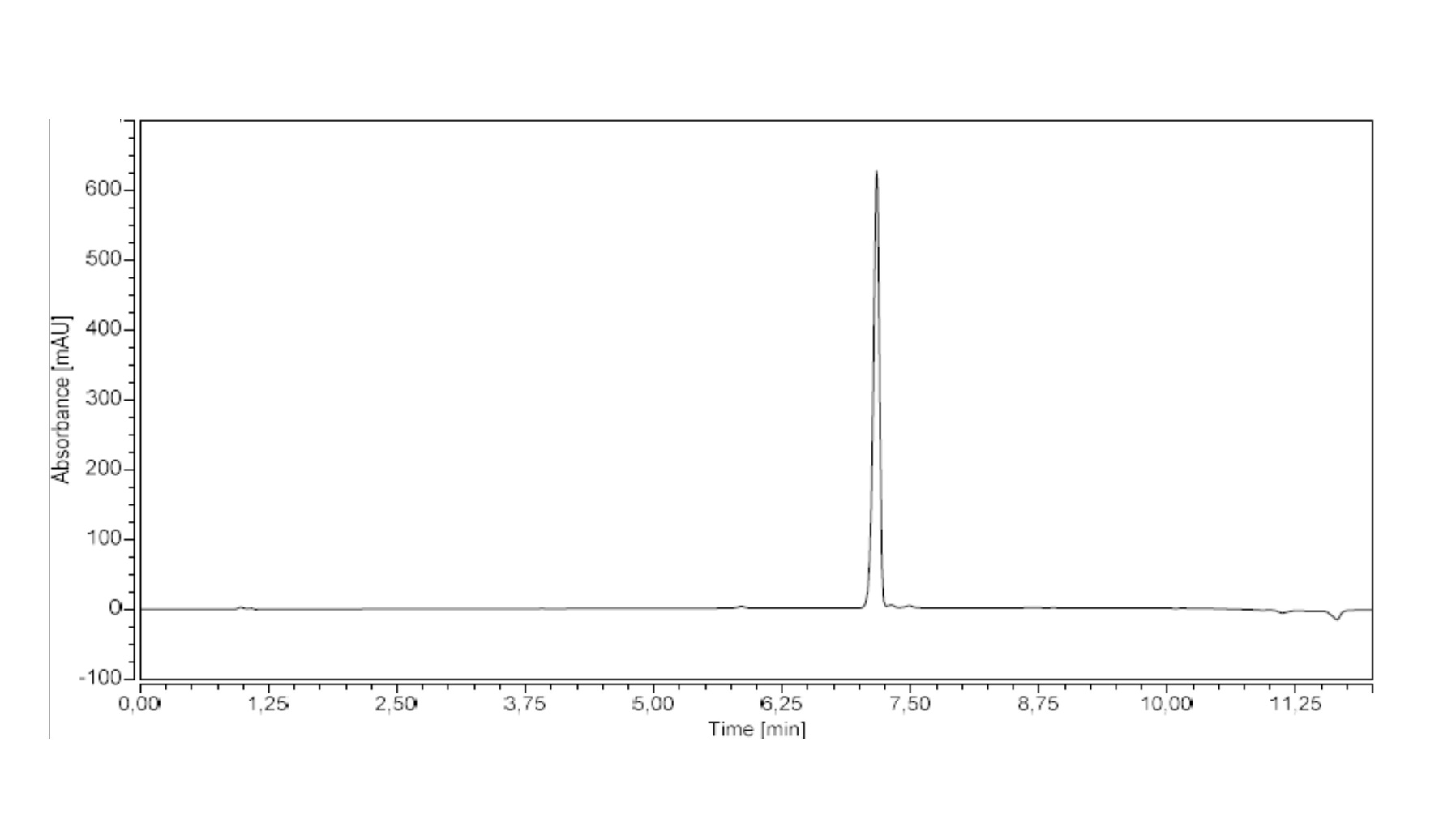

Figure S2D: HPLC purity (210 nm) of RF3458.

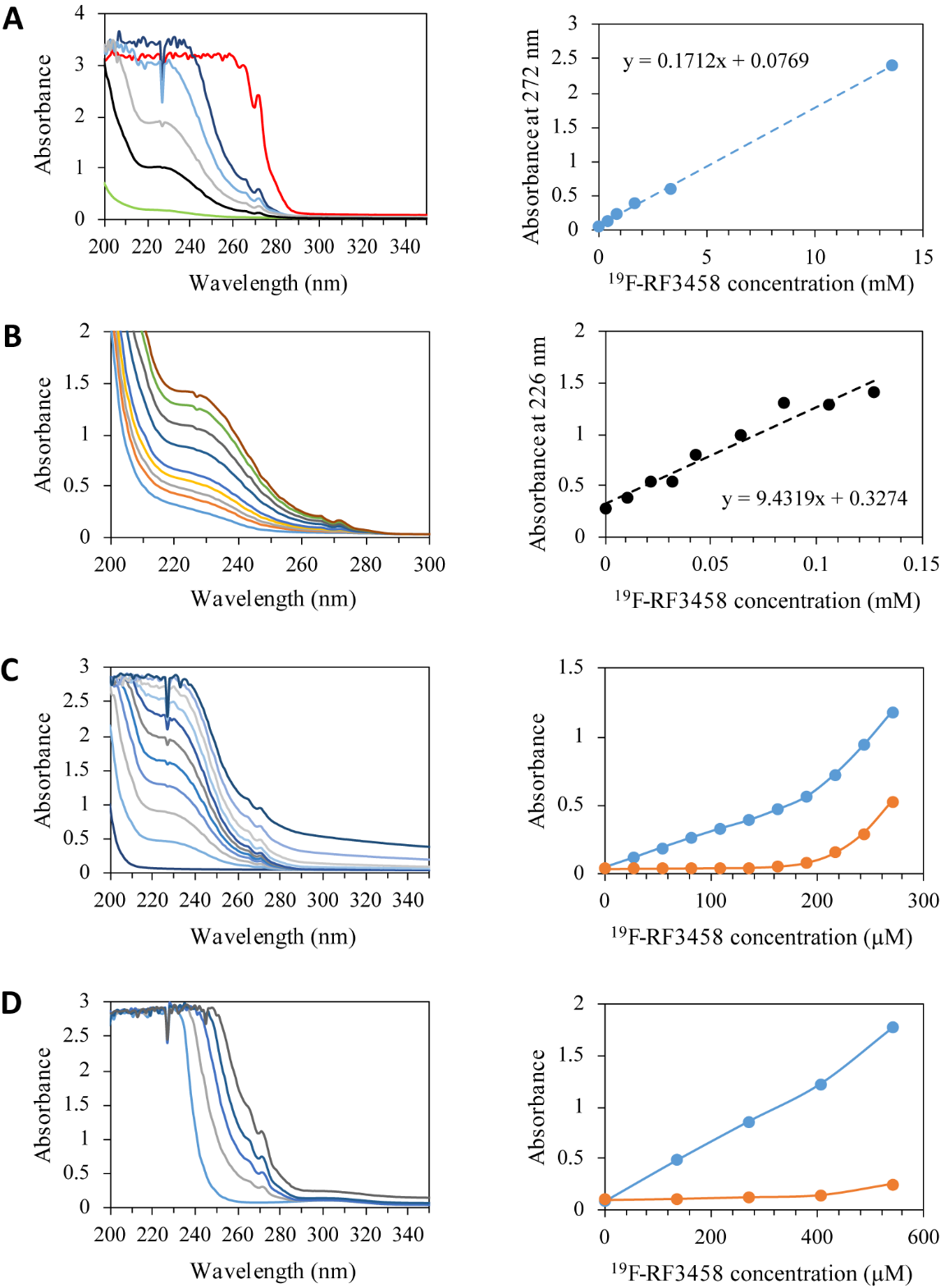

Figure S2: Absorbance spectra of ^19^F-RF3458. (A) Solubility in EtOH (green), pure stock solution (~14 mM in red) was diluted 2, 4, 8 and 16 times (dark blue, light blue, grey and black, respectively in left panel) enabling determination of extinction coefficient at 272 nm (right panel). (B) Increasing amount of ^19^F-RF3458 were added in EtOH (left panel) enabling determination of extinction coefficient at 226 nm (right panel). (C) Solubility in phosphate buffer: increasing amount of ^19^F-RF3458 was added in 40 mM phosphate buffer at pH 6.5 (left panel). Onset of saturation-precipitation is observed above 200 µM (right panel) as revealed by the increase of absorbance at 300 nm (orange) and deviation of linearity at 260 nm (blue). (D) Solubility in phosphate buffer containing 0.1% DPC detergent: increasing amount of ^19^F-RF3458 were added in phosphate buffer at pH 6.5 (left panel). Onset of saturation-precipitation is observed above 400 µM in the presence of 0.1% DPC (right panel) as revealed by the increase of absorbance at 300 nm (orange) and deviation of linearity at 260 nm (blue).
